# Two-way microscale interactions between immigrant bacteria and plant leaf microbiota as revealed by live imaging

**DOI:** 10.1101/695734

**Authors:** Shifra Steinberg, Maor Grinberg, Michael Beitelman, Julianna Peixoto, Tomer Orevi, Nadav Kashtan

## Abstract

The phyllosphere – the aerial parts of plants – is an important microbial habitat that is home to diverse microbial communities. The spatial organization of bacterial cells on leaf surfaces is non-random, and correlates with leaf microscopic features. Yet, the role of microscale interactions between bacterial cells therein is not well understood. Here, we ask how interactions between immigrant bacteria and resident microbiota affect the spatial organization of the combined community. By means of live imaging in a simplified *in vitro* system, we studied the spatial organization, at the micrometer scale, of the bio-control agent *Pseudomonas fluorescens* A506 and the plant pathogen *P. syringae* B728a when introduced to pear and bean leaf microbiota (the corresponding native plants of these strains). We found significant co-localization of immigrant and resident microbial cells at distances of a few micrometers, for both strains. Interestingly, this co-localization was in part due to preferential attachment of microbiota cells near newly formed *P. fluorescens* aggregates. Our results indicate that two-way immigrant bacteria – resident microbiota interactions affect the leaf’s microscale spatial organization, and possibly that of other surface-related microbial communities.

## Introduction

Leaf surfaces constitute a huge microbial habitat that is inhabited by diverse microbial populations including bacteria, yeast, and filamentous fungi (1-9). Bacteria are the most abundant organism among the leaf microbiota (3). The bacterial population of a typical leaf is comprised of hundreds of species of diverse phyla (3, 6, 10). Bacterial cell densities on leaves may reach around 10^5^-10^6^ cells per cm^2^, and cells are observed both as solitary cells and as aggregates that often comprise of several species (3, 11-13). The spatial organization of bacterial cells on the leaf surface is not uniform, and bacterial colonization correlates with the leaf’s microscopic heterogeneity. Bacterial cells and aggregates have been shown to more likely colonize veins, trichome bases, stomata, and the cavities between epidermal cells (1, 8, 11, 14-16). It is thus clear that leaf microscale heterogeneity plays an important role in determining microbial organization on leaf surfaces. However, it is not well-understood whether, how, and at which spatial scales, cell-to-cell interactions affect the microscale spatial organization of leaf microbiota cells.

Mapping the spatial organization of the leaf microbiota population at micrometer and single-cell resolution is a challenge. Thus far, it has been done by means of microscopy of leaf inoculation experiments with a small number of interacting fluorescently-tagged model strains. This approach has revealed non-random dual-species spatial organization at the micrometer scale (15, 17). Other studies analyzed the natural microbiota on leaves using FISH probes (13, 18). These studies highlighted the complex interspecies interactions within natural leaf microbiota and shed light on the spatial scales of interspecies interactions, but were still limited to a low taxonomic resolution. The aforementioned studies suggest that interspecies interactions, at a micrometer spatial scale, are common on leaves. Yet, even if we could accurately map the microscale organization of cells on natural leaves, it would be hard to assess what is the net contribution of cell-to-cell interactions to the observed spatial organization, as leaf surface heterogeneity may mask this information.

Several studies have sought to understand the microscale colonization patterns of immigrant bacteria upon arrival to a new leaf, and how this organization affects cell survival. The fate of individual bacteria was found to depend upon variation in local conditions including e.g., nutrient concentrations and hydration conditions, that modulate the carrying capacity of microsites (19, 20). Moreover, aggregates of resident bacteria have been shown to facilitate the survival of immigrant bacteria cells that join them, on leaf surfaces (21, 22). The majority of these studies were not based on continuous live imaging, therefore they could not resolve whether cells preferentially attached to resident bacterial aggregates, or if cells that randomly attached near or onto aggregates had better survival rates.

Microscale interactions are also important for better understanding early colonization of pathogens and biocontrol agents. Little is known about the microscale interactions of an immigrant pathogen with the resident microbiota and their role in the pathogen’s establishment. Because foliar pathogens typically spend some time as epiphytes before penetrating the leaf interior and causing disease, it is important to understand what they do and how they survive on the leaf surface. In addition, the early colonization of biocontrol agents is also of great interest. As these agents’ establishment on the leaf is desirable, it is of interest to understand how interactions with the native microbiota affect their colonization.

In a previous study, we suggested that preferential attachment of cells to existing aggregates can improve survival in environments exposed to periodic stress (23). That work was based on an individual-based modeling approach that used computer simulations of foraging planktonic cells colonizing a surface under alternating wet-dry cycles. One of the questions we ask here is whether preferential attachment is observed experimentally.

This work examines how environmental leaf microbiota and artificially-introduced immigrant bacteria, spatially organize with relation to each other, post-inoculation. To focus on cell-to-cell interactions and to rule out the impact of leaf surface heterogeneity, we used a simplified experimental system based on glass-bottom multi-well plates. Two model strains were used as immigrant bacteria: *Pseudomonas syringae* B728a and *P. fluorescens* A506 (hereinafter: *Ps* B728a and *Pf* A506). *Ps* B728a is a foliar pathogen model strain that is the cause of brown spot on bean leaves (24-26). *Pf* A506 is a model bio-control agent that is used against *Erwinia amylovora*, the cause of fire blight disease in pears and apples (27, 28). These two strains were introduced to natural microbiota extracted from the surface of fresh leaves. Live imaging microscopy was used to track the microscale spatial organization of both immigrant bacteria and resident microbiota for up to 13 hours post inoculation. Analyses of the spatiotemporal organization of the combined population of the immigrant cells and the resident microbiota on the surface, at micrometer resolution, enabled us to study non-random mutual spatial organization patterns of immigrant cells and native microbiota.

## Materials and Methods

### Experimental design

A description of our experimental design appears in Fig. 1. Briefly, fresh leaves were picked from organically-grown, open-field green bean plants (*Phaseolus vulgaris*) and from an organically-grown pear tree (*Pyrus communis*) (Fig. 1A). The leaf surface microbiota of each plant was extracted as described below (Fig. 1C). In addition, to better represent the chemical environment of the leaf, leaf solutions were prepared for each plant species by blending fresh leaves (Fig. 1D). Fluorescently tagged *Ps* B728a or *Pf* A506 cells (Fig. 1B), which acted as the immigrant bacteria in our experiments, were inoculated into glass-bottom multi-well plates containing leaf solution media, with and without the natural resident microbiota extracted from bean and pear leaves, and surface colonization was tracked through live imaging (Fig. 1E-F).

**Fig. 1.**
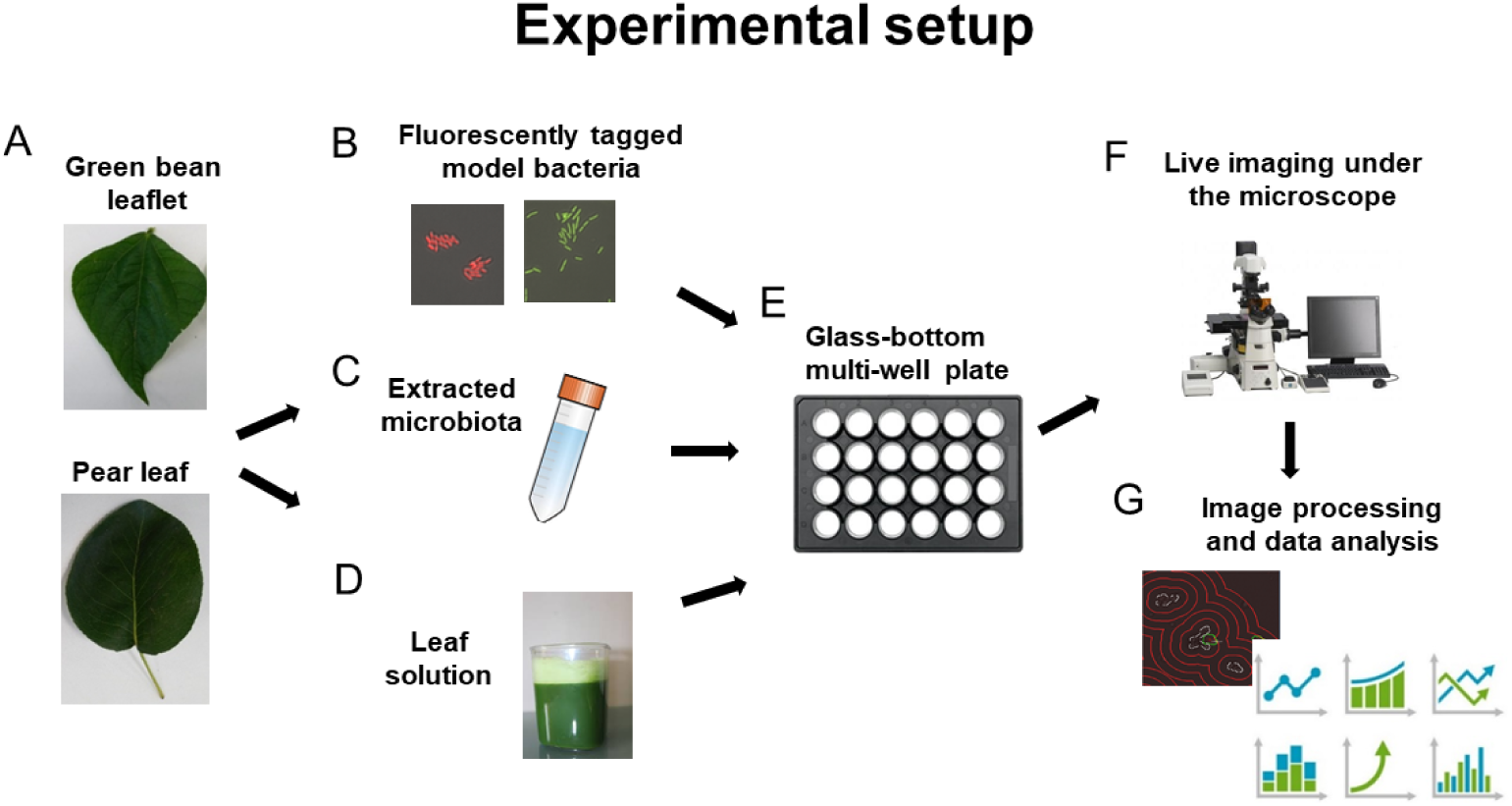
Experimental Setup. **A**. Fresh green bean and pear leaves were picked. **B**. Fluorescently tagged *Ps* B728a and *Pf* A506 were used as immigrant bacteria. **C**. The natural microbiota were extracted from leaf surfaces **D**. Leaf solution was prepared from each plant. **E**. *Ps* B728a or *Pf* A506 were inoculated into glass-bottom wells with leaf solution only, or with both leaf solution and microbiota. **F**. Surface colonization was tracked over time under the microscope. **G**. Image processing and data analysis were performed using custom software in MATLAB and DAIME(32).

### Strains and culture conditions

*Pseudomonas syringae* B728a (25) and *Pseudomonas fluorescens* A506 (28), isolated respectively from green bean and pear leaves, were kindly gifted by Steven E. Lindow, UC Berkeley. *Ps* B728a carried the pKT-TRP plasmid for a constitutive GFP expression (21), whereas *Pf* A506 was transformed using the Tn7 transposon system (29) to produce strains carrying an inserted gene cassette containing a constitutively expressed mCherry gene (this study). Cultures of *Ps* B728a and *Pf* A506 transformants were grown overnight in LB broth media, supplemented with either 30 mg/ml gentamicin or 50 mg/ml kanamycin respectively, under agitation of 400RPM at 28°C. Subsequently, the strains were diluted in sterile phosphate-saline buffer (PBS 1X, 137 mM NaCl, 2.7 mM KCl, 8 mM Na_2_HPO_4_, and 2 mM KH_2_PO_4_) and inoculated into 24-well plates containing leaf solution only or leaf solution supplemented with natural leaf surface microbiota as described below.

### Leaf surface microbiota extraction

Fresh leaves from a pear tree and from green bean plants were sampled from organically grown plants (pear from Kiryat Malakhi area and bean from near Modi’in, Israel) in order to retrieve the microbiota associated with their surfaces. To extract the natural epiphytic microbiota from the leaf surface, we applied a protocol developed in our lab that is based on mechanical scraping. This method preserved most small and mid-size (∼50 µm) aggregates intact, and efficiently recovered the majority of microbiota cells from the leaf surface. This method is an alternative method to the one proposed by Morris et al. that used agitation (30); while we found our method to yield qualitatively similar outcomes (i.e., extracted microbiota), the scraping methods seemed somewhat better at extracting more surface-attached aggregates, in keeping them intact, and in avoiding the possible bias introduced by filtration (see more about a comparison between the two methods in Supp. Fig. S1). Both the abaxial and adaxial sides of the leaves were individually scraped using a disposable cell spreader, after shallowly submerging the leaf in about 10 ml of sterile PBS while ensuring that the petiole was not submerged in the solution. The scraped microbiota was then transferred to sterile 50 ml Falcon tubes, and the remaining loosely attached microbes were re-suspended in about 1-3 ml of PBS from the post-scraping leaf. All leaves utilized in this experiment were picked within 12 hrs of scraping (and an additional 1 hour until microscope screens).

### Leaf solution

Leaf solutions of both pear and green bean leaves were prepared separately by blending 2 to 4 leaves, depending upon leaf size, in 500 ml of PBS 1X. The resulting solution was then autoclaved and filtered through 0.2 µm filters (Millex-GV) to remove leaf particles and intact microbial cells.

### Microscopic analysis of the spatial organization of cells and microbiota on the surface

*Ps* B728a and *Pf A*506 strains were grown overnight in LB broth media to a mid-log phase (OD_600_ = 1.0, 1 cm path length). The culture OD was first adjusted to OD = 0.5, and then further diluted by 1×10^3^ in PBS (by employing serial dilutions). Leaf microbiota extractions (leaf wash) were diluted fivefold (pear) or tenfold (bean) to ensure that the microbiota density was not too high for the required spatial analysis (i.e., to ensure that the surface was not too densely covered by microbiota). Subsequently, 150 µl of the leaf wash product containing the microbiota from either pear or bean leaves was gently inoculated into 24-well plates (24-well glass bottom plate #1.5 high-performance cover glass - Cellvis, USA) containing 800 µl of the corresponding leaf solution (i.e., pear or green bean leaf solution) per well. Experiments were done in triplicates (i.e., 3 wells per each immigrant and microbiota combination). Immediately after microbiota inoculation, the well plates containing the pear or bean systems were then inoculated with 50 µl of diluted *Pf* A506 or *Ps* B728a. 24 well plates (lid on) were mounted on a stage top without warming (room temperature, ∼25°C) during image acquisition.

Images were collected using an Eclipse Ti-E inverted epi-fluorescence microscope (Nikon, Japan) equipped with a Plan Apo 40x/0.95 NA air objective and the Perfect Focus System for maintenance of focus over time. An LED light source (SOLA SE II, Lumencor) was used for fluorescence excitation. *gfp* fluorescence was excited through a 470/40 filter, and emission was collected with a T495lpxr dichroic mirror and 525/50 filter (filter cube #49002, Chroma). mCherry fluorescence was excited through a 560/40 filter and emission was collected with a T585lpxr dichroic mirror and 630/75 filter (filter cube #49008, Chroma). Images were acquired with a SCMOS camera (ZYLA 4.2PLUS, Andor Technology Ltd., UK) controlled with NIS Elements 5.02 software (Nikon Instruments Inc., USA). For time-lapse experiments, images were collected every 30 min. At each time-point, 3×3 adjacent fields of views (covering a total area of 0.98mm^2^) were monitored per each well. Multiple stage positions were collected using a motorized encoded scanning stage (SCANplus IM 130 × 85, Märzhäuse).

### Image processing and segmentation of microbial cells

The segmentation of native microbiota structures and immigrant bacteria cells on the surface was performed in two stages: First, the entire population on the surface was masked by Hybrid Range Filters, which identified objects in the bright field (BF) channel against the background (31). Second, each pixel of the objects was classified as either ‘immigrant bacteria’ or ‘native microbiota’ based on the intensity in both fluorescent channels and a manually calibrated piecewise-linear separator of RED vs GREEN channel intensities. To avoid misclassification, a margin of 1 μm was defined around immigrant bacteria cells (or cell clusters), and any microbiota-labeled pixels in this border zone were removed from the segmentation. As the microbiota extraction method was not 100% free of plant-derived particles such as chloroplasts and tiny pieces of leaf epidermal cells, we further classified the microbiota ‘class’ into plant-derived and microbiota-derived particles. We did this by manual classification, based on examination by eye of the overlay channels. While most plant-derived particles are characterized by their shape and auto-fluorescence, this was difficult to achieve algorithmically by a computer. We therefore decided to do this manually, twice, by two independent persons. By this means, between 0.4%-3.9% of the objects were classified as plant-derived particles (reflecting between 3% to 15% of the total area that was covered by particles on the well bottom). Overall statistics of plant-derived particles are presented in Supp. Table S1, and example images of classifications are provided in Supp. Fig S2.

### Spatial Analysis

#### Colonization pattern of immigrant bacteria in the absence of native micropbiota

Point correlation analysis was carried out by 2D dipole analysis (one-population, between-objects, whole ref. space) using DAIME software (32), and are shown in Supp. Fig. S3. Mean and 95% confidence intervals (CI) were based on analyses of 9 sections (3×3 tiles) of each image (image area is 0.98 mm^2^; one image was analyzed per well).

#### Co-localization of immigrant and microbiota cells or plant parts

Pair Cross-Correlation (PCC) and Nearest Neighbor (NN) analyses were carried out by 2D dipole analysis (two-populations, 500,000 random dipoles) and inflate analysis respectively, using DAIME software (32, 33). Mean and 95% confidence intervals were based on analyses of 9 sections (3×3 tiles) of each image (image area is 0.98 mm^2^; one image was analyzed per well). Note that due to the classification process described earlier, it was necessary to use a narrow margin (of 1 µm) around immigrant cells to avoid misclassification of immigrants as microbiota. Thus, distances up to 1 µm are interpreted as exactly 1 µm. Distance values over 1 µm are unaffected by this artifact.

#### Alterative null random models of spatial organization

In addition to DAIME’s pixel-based randomization null-model, alternative random models that preserve the size and shape of immigrant and microbiota cells and clusters were considered. (1) Object-randomization, representing a random spatial organization, was produced by a Monte Carlo process: The reference population remained unchanged. All discrete objects in the analyzed population were picked in a random order, and each object (preserving its exact size and shape) was assigned a random location with no intersection with previous objects or with the reference population. The randomized datasets were analyzed using DAIME in a similar manner as the real data. Graphs in the relevant supplementary figures show the mean and 95% confidence interval of five different randomizations for each of the nine tiles (45 in total, per well). (2) label-randomized datasets, where microbiota and immigrant bacterial structures’ labels were randomly permutated. Graphs in the relevant supplementary figures show the mean and 95% confidence interval of 100 different randomizations.

#### Aggregate-size distributions

In order to cluster tightly localized cells as aggregates, we performed morphological closing operations on the segmented masks with a structuring element of 2 μm radius.

## Results

We first performed surface colonization experiments with *Ps* B728a or *Pf* A506 cells in leaf solution only (i.e., without leaf microbiota cells, Fig. 1). Inoculation with bean leaf solution led to a more rapid and intense surface colonization of both strains than that with pear leaf solution (Fig. 2), possibly due to higher nutrient concentrations (consistent with growth curves in leaf solution in bulk, Supp. Fig. S4). *Pf* A506 cells tend to form aggregates on the surface, which grow over time. Typically, a nascent *Pf* A506 aggregate begins with the attachment of a single cell, and most of its progeny remains in the same location (see Supp. Figs. S5, S6). *Ps* B728a cells typically do not form pure aggregates, but rather colonize the surface more homogenously (see Supp. Figs. S5, S6). To test if the surface colonization pattern, with leaf solution but in the absence of microbiota, deviates from expected from random colonization, we performed pair-correlation spatial analyses (see Materials and Methods). These analyses showed that the spatial distribution of both *Pf* A506 and *Ps* B728a cells (or cell clusters) do not considerably deviate from the expected by chance (with one minor exception - see Supp. Fig. S3). These results also indicate that the surface is homogenous, and that there are no artifacts such as tiny submicron leaf pieces in the leaf solutions that might have attracted cells.

**Fig. 2.**
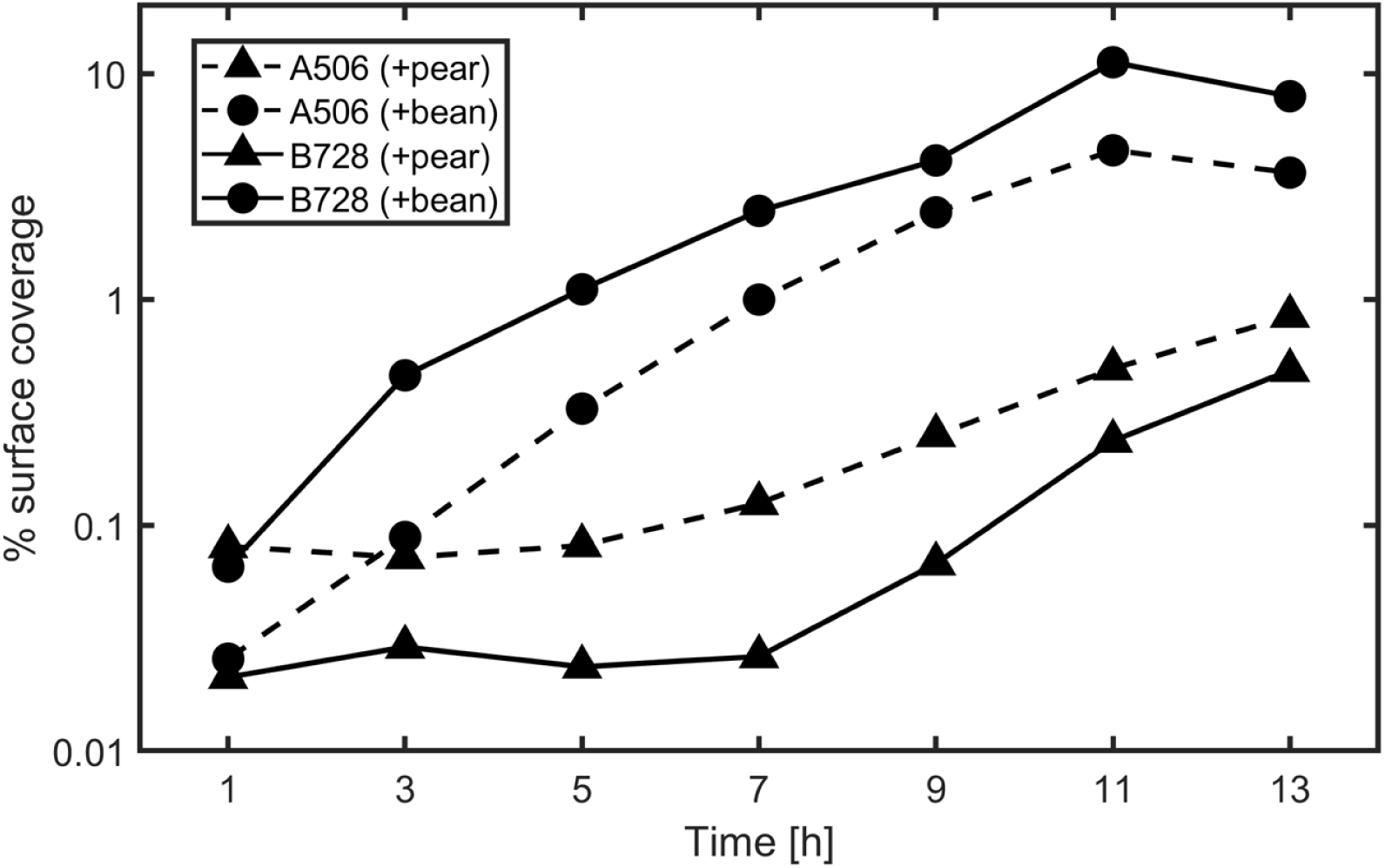
Surface colonization dynamics without microbiota (leaf solution only). The graphs show the percentage of well surface covered by cells as a function of time.

Next, we moved on to experiments with both immigrant and resident microbiota. The leaf microbiota extracted from both plants included bacteria, yeast-like cells, and fungi in both solitary and aggregated forms (as observed by microscopy, Fig. 3). Many of the aggregates were ‘mixed’, i.e., comprised of both bacteria and yeast cells. Microbiota of both leaves contained mostly smaller aggregates and fewer large ones of up to a few hundred µm^2^ (Fig. 3 and Supp. Fig. S7 E-H). To study the spatial organization of the combined population of immigrant and resident leaf microbiota, we tracked the respective surface colonization over time under the microscope (every 30 minutes for 13 hours). At each time point, we identified immigrant cells and all microbiota components through image analysis (Materials and Methods). Particles identified as plant derived were excluded from the analysis (Materials and Methods). As time passed from the inoculation point, an increasing number of both immigrant and microbiota cells were observed on the well bottom surface (Fig. 4, Supp. Fig. S7, S8-S11). Immigrant cells and microbiota cells were observed attaching to the surface, moving, dividing, detaching, and some of them forming cell clusters (micro-colonies) in the form of surface-attached aggregates. Of the microbiota cells, aggregates were observed sinking to the bottom surface (most aggregates sank over the first 2-3 hours, as might indicated from the temporal changes in aggregate-size distributions, Fig. S7 E-H), while some solitary cells were observed too, some of them clearly motile.

**Fig. 3.**
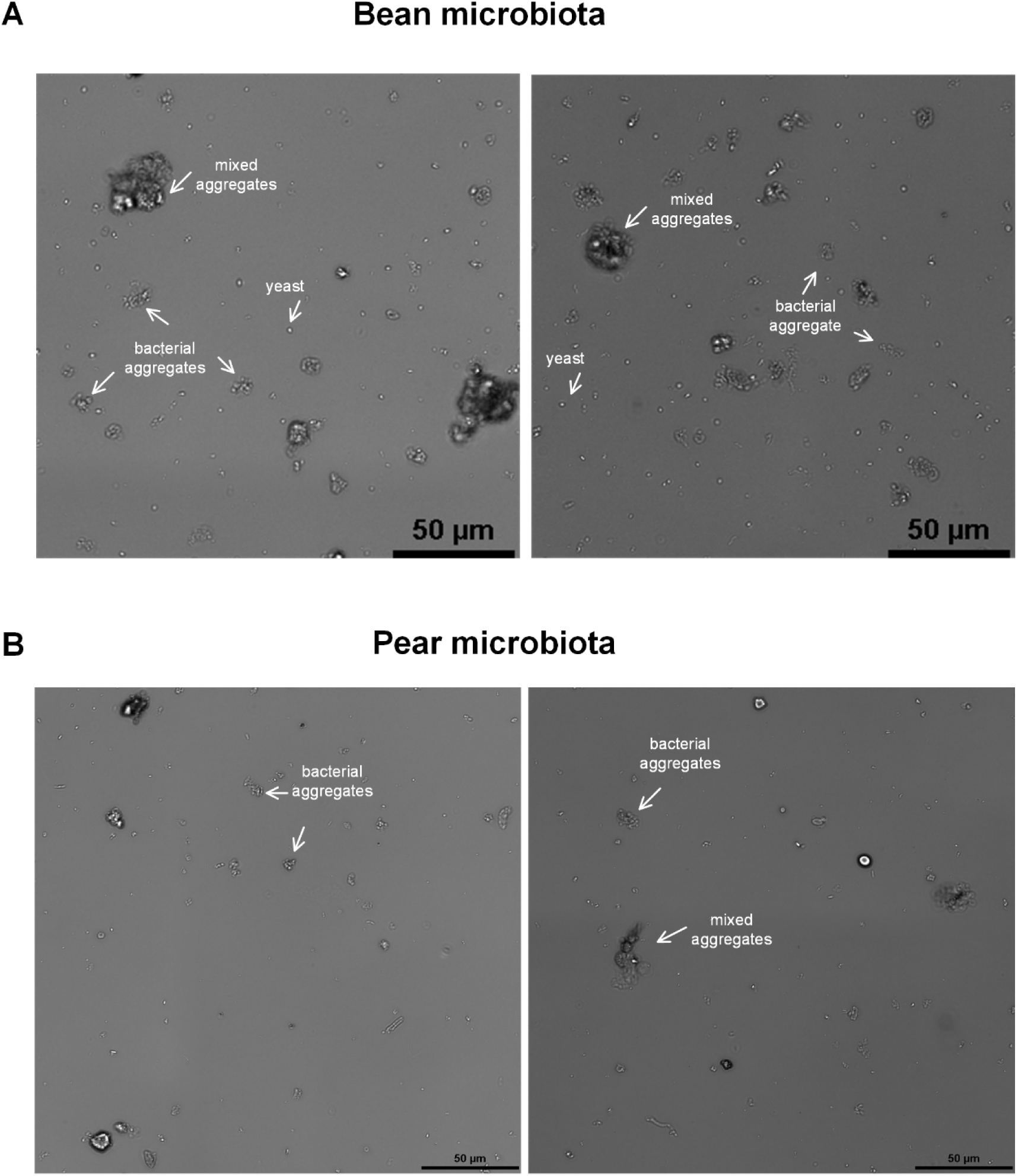
Extracted microbiota from bean and pear leaves. Two representative images of the extracted microbiota from bean (**A**) and pear (**B**) leaf surfaces, as observed on glass-bottom well plates, prior to immigrant bacteria colonization.

**Fig. 4.**
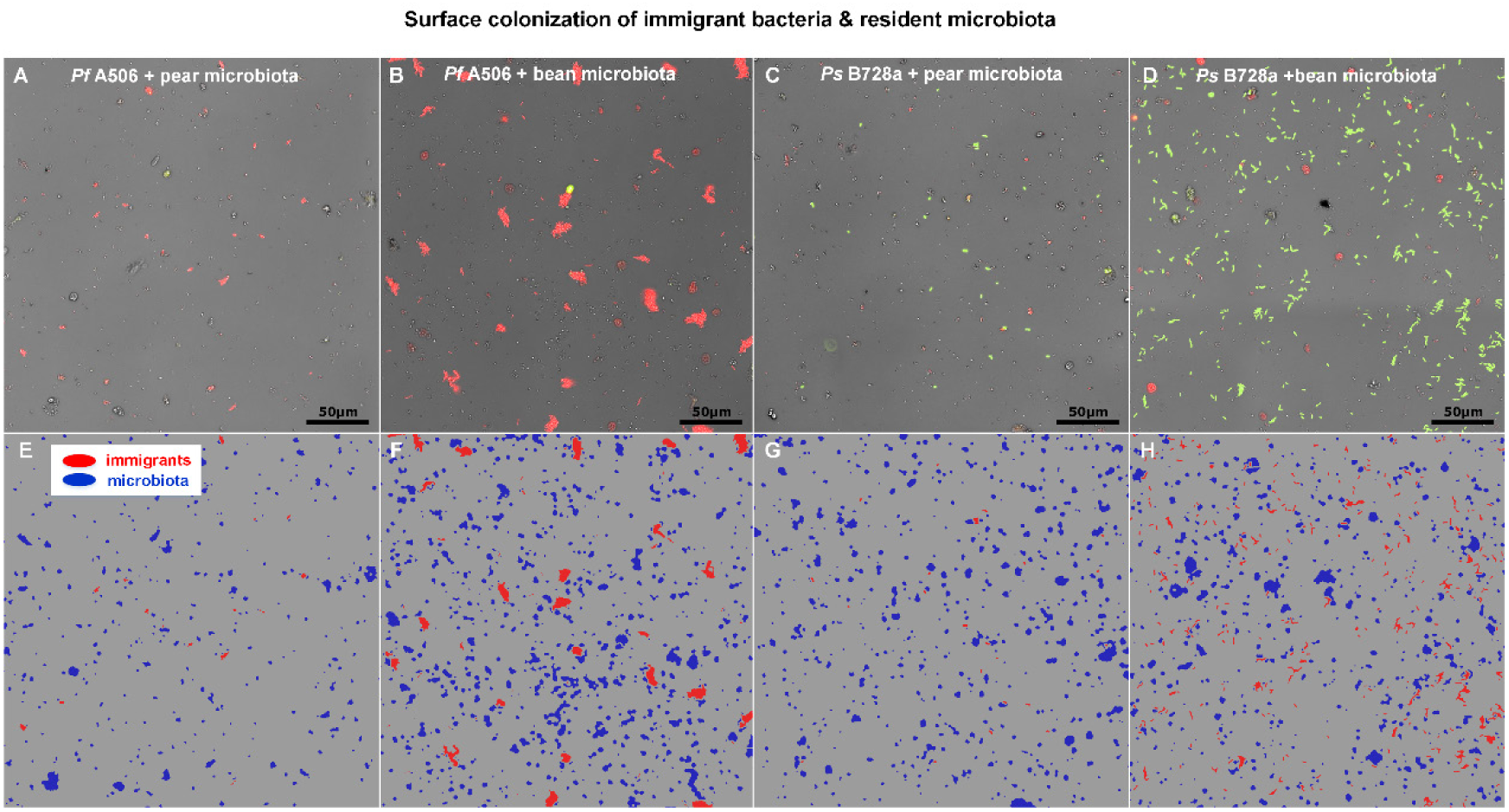
Surface colonization of immigrant bacteria introduced to resident microbiota. **A**. *Pf* A506 (red) with pear microbiota. **B**. *Pf* A506 (red) with bean microbiota. **C**. *Ps* B728a (green) with pear microbiota. **D**. *Ps* B728a (green) with bean microbiota. All images were taken 8 hrs post inoculation. Note the differing colonization patterns of the two strains: *Pf* A506 formed aggregates, while *Ps* B728a remained mostly as solitary cells that colonized the surface in a less ‘clustered’ pattern (see also Supp. Fig. S7). **E-H**. The corresponding classification of the overall population into immigrant cells (red) and microbiota components (blue). Objects suspected as plant particles were removed.

Next, we wanted to know whether and how the spatial organization of resident microbiota cells and immigrant cells on the surface was affected by their co-existence. More specifically we asked: Do immigrant cells and microbiota cells co-localize at the microscale more than expected by chance? To answer this question, we employed two commonly used, complementary methods for spatial analyses of our spatial data of the microscale organization of both immigrant and microbiota components on the well surface (Fig. 4 E-H). We used DAIME, a common tool for image and spatial analysis in microbial ecology (32), which implements both methods. The first method, termed pair cross-correlation (PCC), used DAIME 2D dipole algorithm (32-34); and the second method, termed nearest-neighbor (NN), used DAIME 2D inflate algorithm (32, 33) (See Materials and Methods).

The overall picture, emerging from the analyses of the experiments of both immigrant strains, was that there were significantly more immigrant cells (or cell clusters) within a small distance, of a few micrometers, from resident microbiota components, than expected by chance (Fig. 5). The extent of deviation from random, and the spatial distances at which this deviation was observed, varied with method type, and between the four combinations of immigrant strain and plant species. Overall, qualitatively similar results were observed for three experimental replicates of each immigrant-microbiota pair (Fig. S12).

**Fig. 5.**
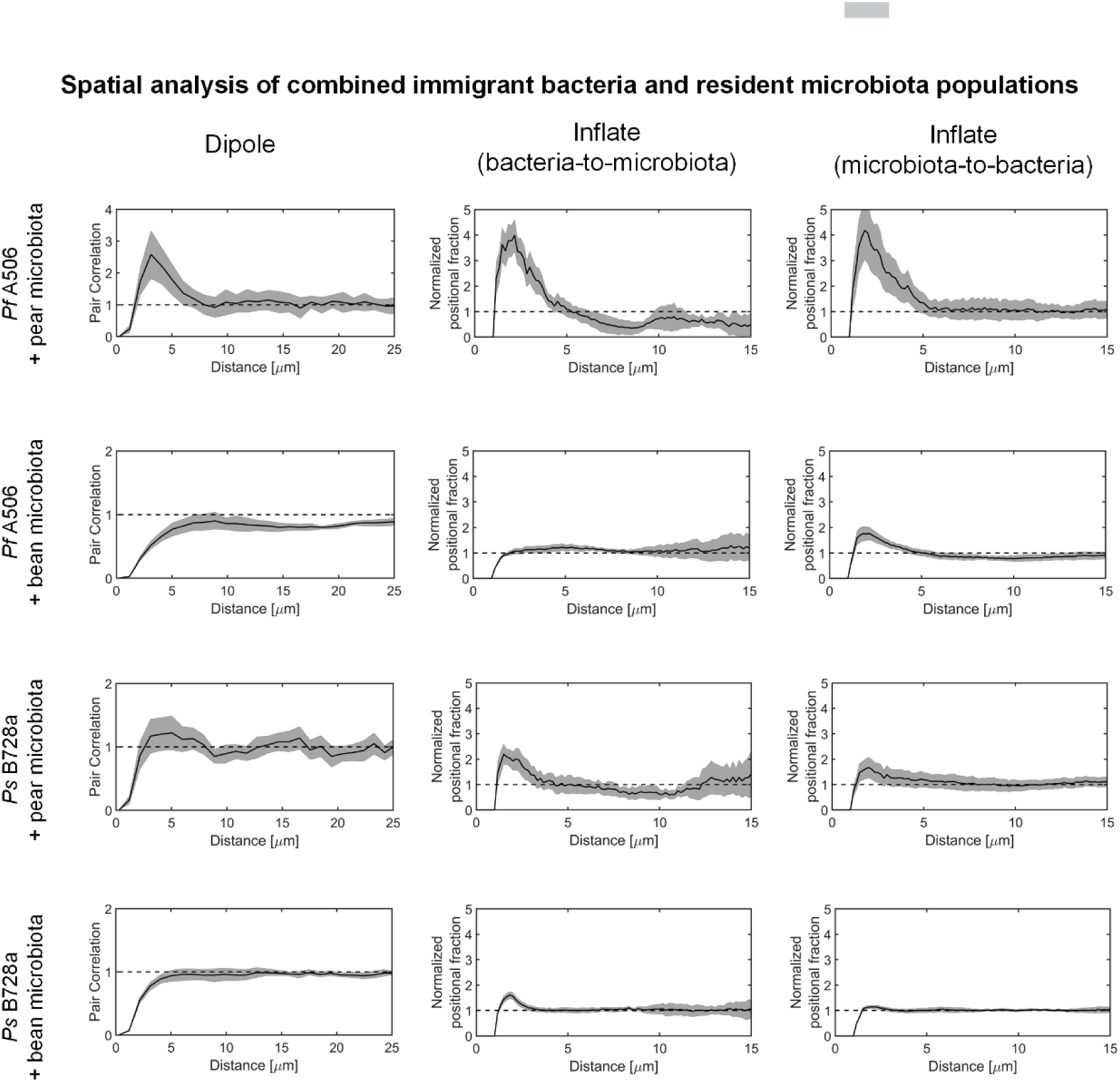
Spatial analysis of combined immigrant bacteria and resident microbiota based on Pair Cross-correlation and Nearest Neighbor methods. Shown are results of pair cross-correlation (PCC), using DAIME 2D dipole algorithm; and nearest-neighbor (NN), using DAIME 2D inflate algorithm (32, 33) (See Materials and Methods). Black line and gray envelopes present mean and 95% confidence intervals. In this type of analysis, deviation from the expected by chance are at distances significantly above or below a value of one. For all combinations of immigrant strain and plant microbiota, at least by one of the methods, there are significantly more cells at short distances to microbiota cells, than expected by chance. Note that when considering an alternative object-based randomization scheme (See Methods), the Dipole algorithm shows co-localization at distances of up to 5-10 µm (Supp. Fig. S15). Note that all minimal observed distances < 1 µm are considered a distance of exactly 1µm (due to technical reasons, see Materials and Methods).

At t = 8 hrs post inoculation, there were significantly more *Pf* A506 cells (or clusters) than expected by chance at distances < ≈5 μm from resident pear microbiota, as reflected by both PCC and NN algorithms (Fig. 5, 1^st^ row). In this type of analysis, at each given distance, values above 1 indicate co-localization of cells (immigrant and microbiota cells co-localize at this distance more than expected by chance) and values below 1 indicate that such co-localizations are less than expected by chance). In the presence of bean microbiota, the *Pf* A506 did not show deviation from random by PCC, but did show a clear deviation in the microbiota-to-immigrant NN analysis (Fig. 5, 2^nd^ row). Although *Ps* B728a showed no deviation from random based on PCC, it showed a clear deviation from random in the NN analyses when co-cultured with both pear and bean microbiota (Fig. 5, 3^rd^ and 4^th^ rows). Co-localization of *Ps* B728a to microbiota components was observed at distance < ≈2-3 μm (Fig. 5, 3^rd^ and 4^th^ rows). The differences in outcome of the PCC versus NN algorithms reflect differences in each algorithm’s respective ability to detect existing spatial patterns and deviations from randomness of various datasets (see more in Discussion).

We further compared the results to those expected by chance based on several alternative null-models. In addition to the pixel-based null-model provided by DAIME, we performed randomization that preserves the size and shape of all immigrant cell clusters and microbiota entities detected on the surface, using Monte Carlo simulations (see Methods). We did it using two approaches: One that randomizes the localization of all immigrant cells clusters with and without allowing overlaps between objects (Fig. S13); and a second approach that randomly permutes the objects’ labels (i.e., its classification as immigrant or microbiota object) (Fig. S14) (see Methods). Using these two stricter randomization models that take into account biologically relevant constraints that (a) preserve the size and shapes of microbial objects and (b) restrict spatial overlap between objects, increased the significance of co-localization at small distances, especially of the Dipole PCC algorithm (Fig. S14, S15).

Finally, we further asked if the observed co-localizations were due to one-way or two-way interactions. The asymmetry of the NN results, when comparing the two directions of the analysis, i.e., for which population the distances to nearest neighbor are compute (immigrant cells or microbiota; second and third columns in Fig. 5), led us to postulate that it tells us something about the underlying process that led to the observed co-localization. For example, whether these co-localizations resulted from preferential attachment of immigrant cells near microbiota cells, or vice versa. To further study this asymmetry, we used our live-imaging data to track all neighboring pairs of microbiota and immigrant bacteria (at distances of up to 2 μm), and checked which element of the pair was present earlier and which one joined subsequently. Interestingly, we found that for *Pf* A506, about 50% of co-localized pairs were a result of preferential attachment of resident microbiota cells to newly formed *Pf* A506 aggregates (Fig. 6A). This was also observed in time-lapse analysis, where it was easy to observe many cases of individual microbiota cells joining newly formed aggregates of *Pf* A506 (Fig. 6B, Supp. Files. S1, S2). This phenomenon is nicely reflected in the asymmetry of the NN analysis, in particular with *Pf* A506 and bean microbiota, where the microbiota-to-immigrant direction of the analysis showed a much stronger signal (reflecting solitary microbiota cells joining large *Pf* A506 aggregates) (Fig. 5). In contrast to *Pf* A506 cells, *Ps* B728a cells were not forming aggregates, and preferential attachment of microbiota cells to *Ps* B728a cells or cell clusters was not observed for either the native or non-native plant microbiota (at least not within the first 9 hrs; Fig. 6A,B, Supp. Files S3,S4). Again, it was nicely reflected in the slightly stronger signal in the immigrant-to-microbiota direction of the NN analysis (Fig. 5).

**Fig 6.**
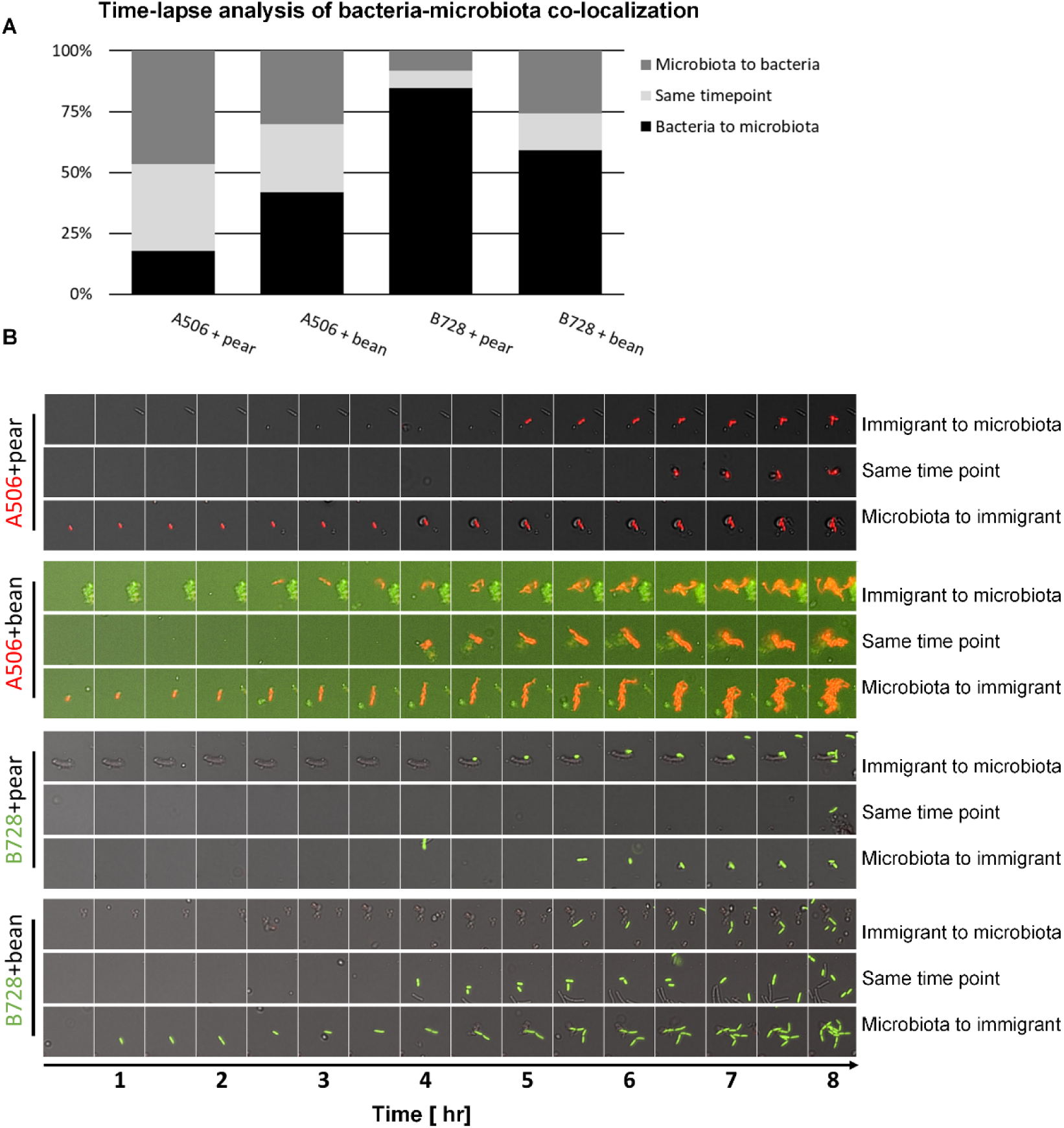
Two-way microscale interaction dynamics. Inspection of co-localized immigrant cells and microbiota shows incidences of all three possible scenarios: immigrant cells joining microbiota cells, microbiota cells joining immigrant cells, and both arriving at the same time point (same 30-minute time window) to the surface. **A**. The fraction of each of the three scenarios per each strain-microbiota pair (based on manual inspection of more than 100 randomly selected co-localized objects). **B**. Example time-series images of immigrant-to-microbiota attachments, same time-point, and microbiota-to-immigrant events (more time series are provided as Supp. Files 1-4).

## Discussion

In this work, we studied the spatial surface colonization pattern of cells of a co-culture of immigrant bacteria and natural resident leaf microbiota cells, through live imaging and spatial analyses at the microscale. We found that both the biocontrol agent *Pf* A506 and the plant pathogen *Ps* B728a display a significant co-localization pattern in co-cultures of either native or non-native resident microbiota (pear and bean). *Pf* A506 co-localization was in part due to attachment of microbiota cells near newly formed *Pf* A506 aggregates, and *Ps* B728a showed a significant co-localization signal without such clear reciprocal preferential attachment.

The non-random co-localization of *Pf* A506 and *Ps* B728a with leaf microbiota, while not necessarily observable by the naked eye, was revealed by a rigorous image and spatial analysis. As our simplified system excludes the heterogeneity of natural leaf surfaces, this non-random organization must originate from interactions between the immigrant and resident microbiota cells. We found that native microbiota and immigrant bacterial cells may exhibit a spatial association, driven by biased attachment and movement on the surface. Our analyses indicate that most such interactions occur on the surface, and not in the bulk fluid: Most microbiota cells settle on the well surface within 2-3 hours, and only 10% to 30% of co-localized immigrant-microbiota components arrived at the same time to the surface (Fig. 6 A). Indeed, detachment, reattachment, and movement of cells were previously suggested to play a role in the spatial organization of cells on leaf surfaces (35).

The spatial scales of interactions are of major importance in microbial ecology (13, 15, 31, 36-38). Our work indicates that the effective scale for such co-localization is at distances of up to ≈5 μm, consistent with the conclusions of previous work on bacterial interactions on leaf surfaces (13, 15). In all four immigrant-microbiota combinations, about 50% (45-56% at 8 hrs) of immigrant cells on the surface were at distances of up to 5 μm from microbiota cells. Co-localization at distances above ≈5 μm between immigrant and resident bacterial cells does not significantly deviate from random (Fig. 5) with the exception of a few cases of less-than-expected co-localizations (i.e., values below 1) at distances of 5-20 μm (Fig. 5, Fig. S15), possibly due to movement of cells within this distance range toward their counterparts.

We note that the various spatial analysis methods have different strengths and weaknesses, and that the differences in the detected deviation from random patterns between methods can be informative. For example, non-random patterns can be masked in PCC, if, for example, there is a big difference in the typical object size between the two populations, and distances are within the same order of scale (as can be seen in Supp. Fig. S3), which may explain the lack of deviation from random in PCC, while such deviation was detected by the NN algorithm. NN is limited in its ability to detect spatial patterns that are characterized by distances that are longer than the typical shortest distances between objects. As demonstrated in our results, the NN algorithm’s asymmetry of results is also of interest, as it may reflect directionality or bias in the underlying process that generates the spatial organization (like the two-way interactions in our experiments).

The choice of a proper meaningful random model for assessing spatial patterns, that takes into account the system’s inherent properties and constraints is important. In particular, we suggest that the object-randomization model that we used, which preserves the size and shape of microbial objects (cells, clusters and aggregates) is important for the analysis of spatial data representing relatively sparse, non-continuously populated spaces (e.g., bacterial colonization on water-unsaturated surfaces as opposed to saturated surfaces with continuous biofilms) and for uncovering patterns that are within the range of the objects’ sizes, and of irregular shapes.

When examining the co-localization of immigrant bacteria and microbiota, comparison with object-based random models increased, in most cases, the significance of co-localization, particularly for the PCC 2D Dipole algorithm. This increase is mainly because microbial ‘objects’ sizes and irregular shapes add a constraint to the randomization – by prohibiting overlaps between objects – which is reflected in the apparent observed repulsion at distances less than, or comparable to, their size (see Supp. Fig. S15).

We note that our experiments were performed with leaf solutions that do not perfectly mimic the chemical environment on the leaf surface. Similarly to what bacteria face on natural leaf surfaces, the leaf solutions were nutrient poor, and contain compounds that microbes encounter on the leaf surface. They do, however, contain other compounds that originate from the leaf internals, including for example, vacuole components. Autoclaving the leaf solution could have also changed some of the solution properties. Because simple leaf washes (from leaf surface only) did not lead to the surface attachment of both studied bacterial strains, we made this compromise in order to study co-localization of immigrant and natural resident leaf microbiota in controlled, more tractable environments. In addition, we note that the extracted leaf microbiota include some plant-derived particles such as chloroplasts and tiny pieces of epidermis. However, as these were only small fraction of all microbiota components (by number and area), we believe that it did not have a major impact on the revealed observed two-way interactions. Our results were robust to the inclusion or removal of these from our analysis (data not shown). Interestingly, some indication that points to co-localization of *Ps* B728a to plant particles at distances of a few μm was also observed (Supp. Fig. S16).

In a previous study, we used modeling and computer simulations to show that preferential attachment can be an efficient general surface colonization strategy for bacteria on ‘patchily colonized’ surfaces that are exposed to periodic stress such as desiccation, antibiotics, and predation (23). The present study provides experimental evidence for preferential attachment during surface colonization. This finding supports previous indications of preferential attachment in an *in vitro* experimental system using a single bacterial strain (39) or environmental microbiota (40). It is likely that biased surface colonization mediated by preferential attachment is a common bacterial organization trait within the phyllosphere, and possibly within other surface-related microbial habitats.

A non-random spatial organization such as the one that we observed, can be a result of various mechanisms. One example is the reported biased surface movement toward dense areas controlled by sticky polysaccharide trails (39). Differential hydrophobic cell-surface and cell-cell adhesion forces could also be at play (41). Other mechanisms may involve chemotactic movement toward aggregates (42), which may underlie the observed gravitation of *Ps* B728a toward microbiota aggregates; or informed attachment decisions, which may be the case in the observed association of *Pf* A506’s and microbiota aggregates. Moreover, intra- and interspecies quorum sensing are possibly involved in such mechanisms (42, 43), especially as quorum-sensing signals on unsaturated surfaces such as plant leaf surfaces are highly localized, and quorum size can be surprisingly small, even as low as a few dozen cells (42, 44). Other sensing systems, such as peptidoglycan sensing by bacteria, can serve as indicators of cell proximity (45).

Regardless of the exact underlying mechanism, spatial organization may confer fitness advantage in environments that select for collective protection from stresses (23). Aggregates have been shown to increase bacterial cell survival on leaves under dry conditions (21, 46). In addition, aggregation has been found to increase survival on drying surfaces under moderate humidity through the formation of larger microscopic droplets around aggregates (47). These microdroplets have been shown to protect cells from desiccation (47). The association of immigrant bacteria and natural microbiota, while not necessarily mutually beneficial, may lead to increased protection. Finally, increased aggregate size may provide better protection from the various stresses that cells experience on leaf surfaces, including desiccation, UV radiation, antibiotics, and predation.

It is not clear if the association that we observed between immigrant cells and microbiota is specific to species or strains of the microbiota. As the leaf microbiota is comprised of hundreds of species, it is reasonable to assume that aggregates’ specific species compositions affect spatial organization. Bacterial cells are known to sense their local environments, and cell colonization and behavior has been shown to be affected by interspecies signaling (48). Initial evidence for such interactions at the microscale, even between closely related strains within the same species, has been recently observed in isolated strains from the rhizosphere (49).

To conclude, our study demonstrates that inter-species microscale interactions likely play a role in the spatial organization of leaf microbiota. In particular, immigrant bacterial cells and resident microbiota tend to co-localize at the microscale, and this co-localization results from two-way interactions between immigrant and resident cells. These findings underscore the importance of individual-based approaches for improving of our understanding of leaf microbial colonization processes, with clear implications for plant pathology and the design of biocontrol approaches. More broadly, our results might be relevant to other surface-associated microbial habitats, including plant and animal microbiomes.

## Acknowledgements

We thank Y. Hadar for valuable comments on the manuscript. We thank S. Lindow for kindly providing bacterial strains, and R. Feuchtwanger from Gan Hasadeh and N. Shachar for providing fresh leaves for this study. J. P. acknowledges the Lady Davis Trust for a postdoctoral fellowship. M. B. acknowledges the Rudin MSc scholarship. This work was supported by a research grant to N. K. from the James S. McDonnell Foundation (Studying Complex Systems Scholar Award, Grant #220020475) and from the Israel Science Foundation (ISF #1396/19).

