## Supplementary Materials for "Two-way microscale interactions between immigrant bacteria and plant leaf microbiota as revealed by live imaging"

Steinberg, Grinberg et. al.

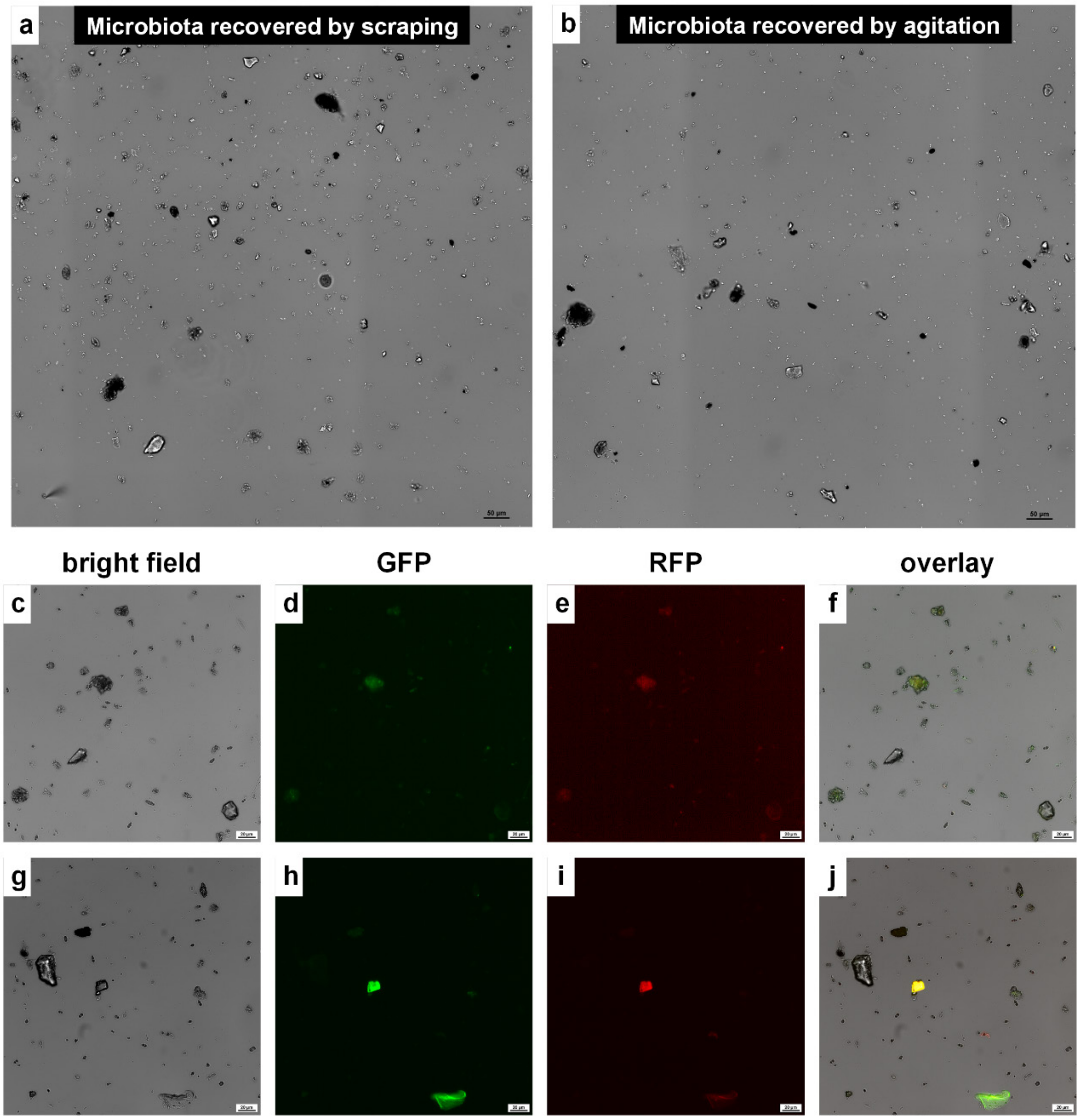


**Fig. S1.** Leaf scraping and leaf agitation methodologies for harvesting leaf microbiota. (a-b) representative images showing the composition of green bean leaf wash extracted by scraping (a) and agitation (b) techniques applicable to recover the natural leaf microbiota. Both methodologies were found to be adequate for harvesting of leaf microbiota including intact bacterial aggregates. Images show a 1x1 mm section on the surface of a glass-bottom well plate. (c-j) Split channel view of crude leaf wash extracted by scraping (c-f) and agitation (g-j). Both scraping and agitation leaf wash includes auto fluorescence particles which in most cases, originate from fractured leaf parts. Images captured with a 40X objective in bright field, GFP channel (Ex 470/40 nm Em 525/50 nm) and RFP channel (Ex 560/40 nm Em 630/75 nm).

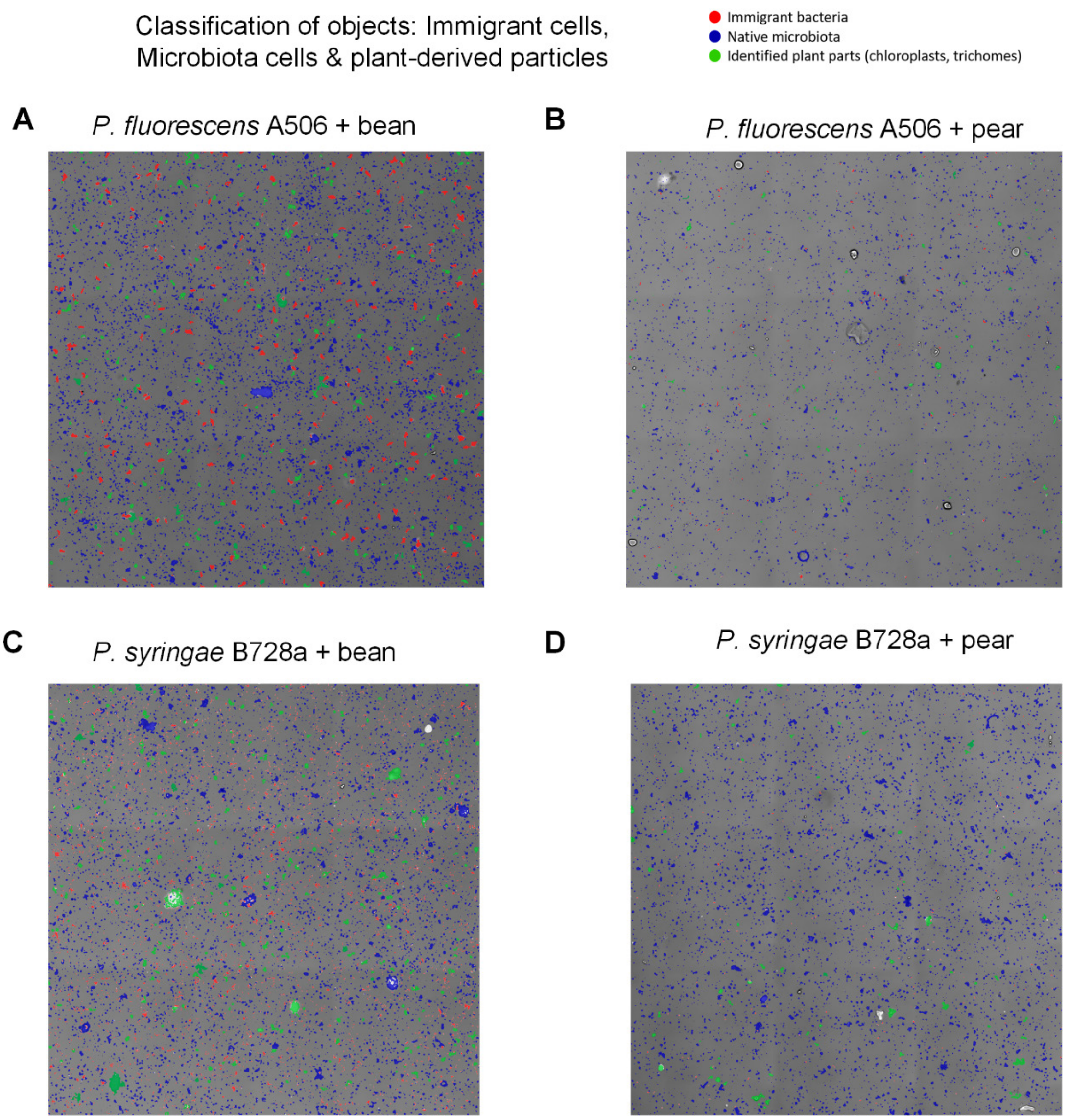


**Fig. S2 Classification of objects to either microbiota or plant derived particles.** Classification was done in two stages. First, the fluorescently-tagged immigrant strains were automatically classified by their GFP and RFP fluorescence intensities. Then, the remaining elements were manually classified as either microbiota or plant particles, twice, each time by a different co-author. The two manual classifications were analyzed, and yielded qualitatively similar results.

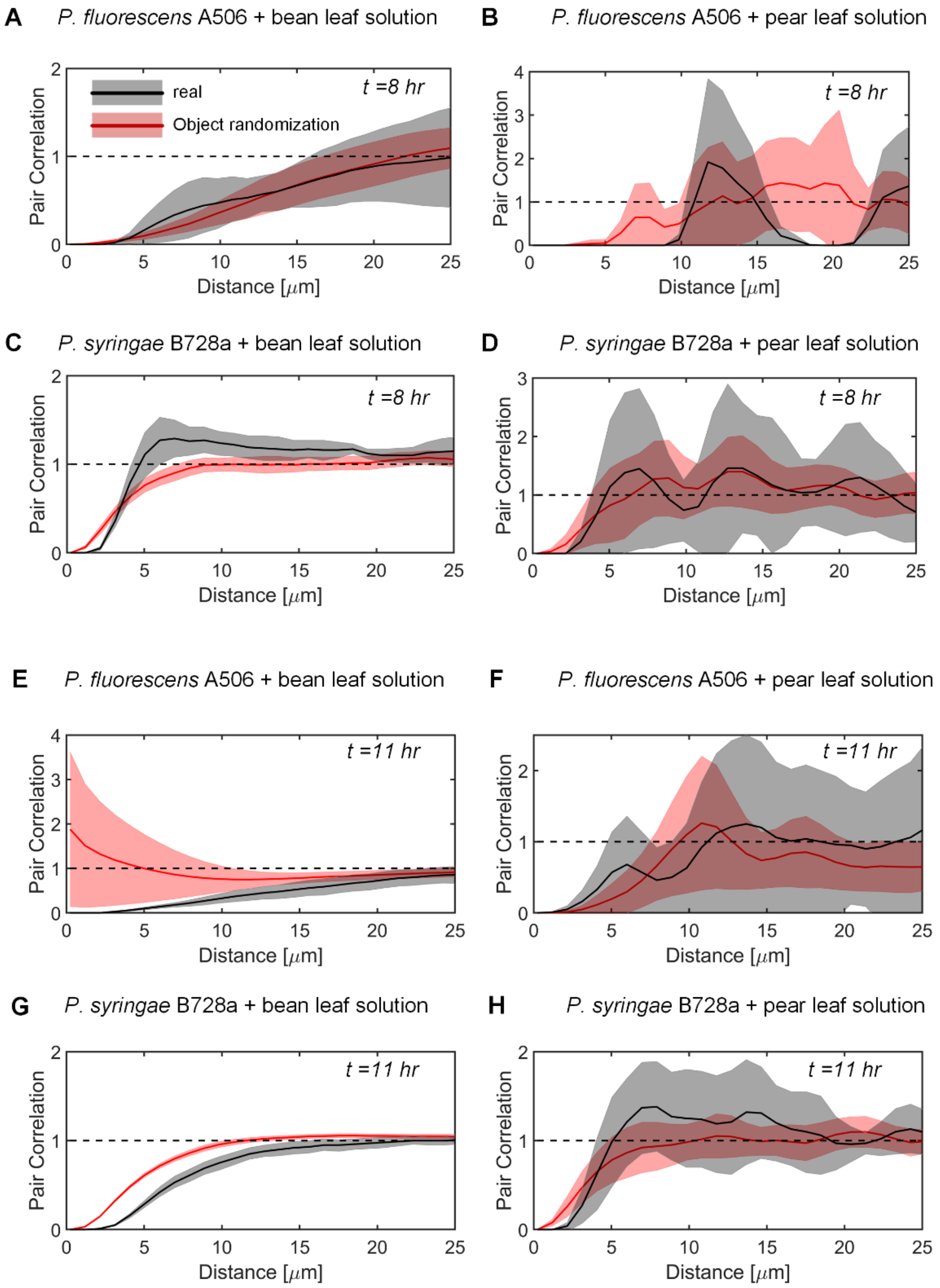


**Fig. S3 Spatial organization of immigrant bacteria with leaf solution.** Pair correlation function of bacterial clusters in leaf solution only, using DAIME 2D dipole analysis. For all combinations of bacteria and leaf solution, the observed pair correlation (in black) 95% confidence interval (in gray), deviates from CSR (dashed line) in small distances, comparable to object sizes. However, a more realistic null model (in red) generated by a Monte Carlo simulation that prohibits overlaps between objects, as described in the Methods, also mostly deviates from CSR, since CSR does not account for the minimal distances imposed by non-negligible object sizes. The confidence intervals of Monte Carlo simulations demonstrate that most of the observed deviation from randomness may be explained by geometric constraints rather than biological processes **A-D** 8 hrs post inoculation. **E-H** 11 hrs post inoculation.

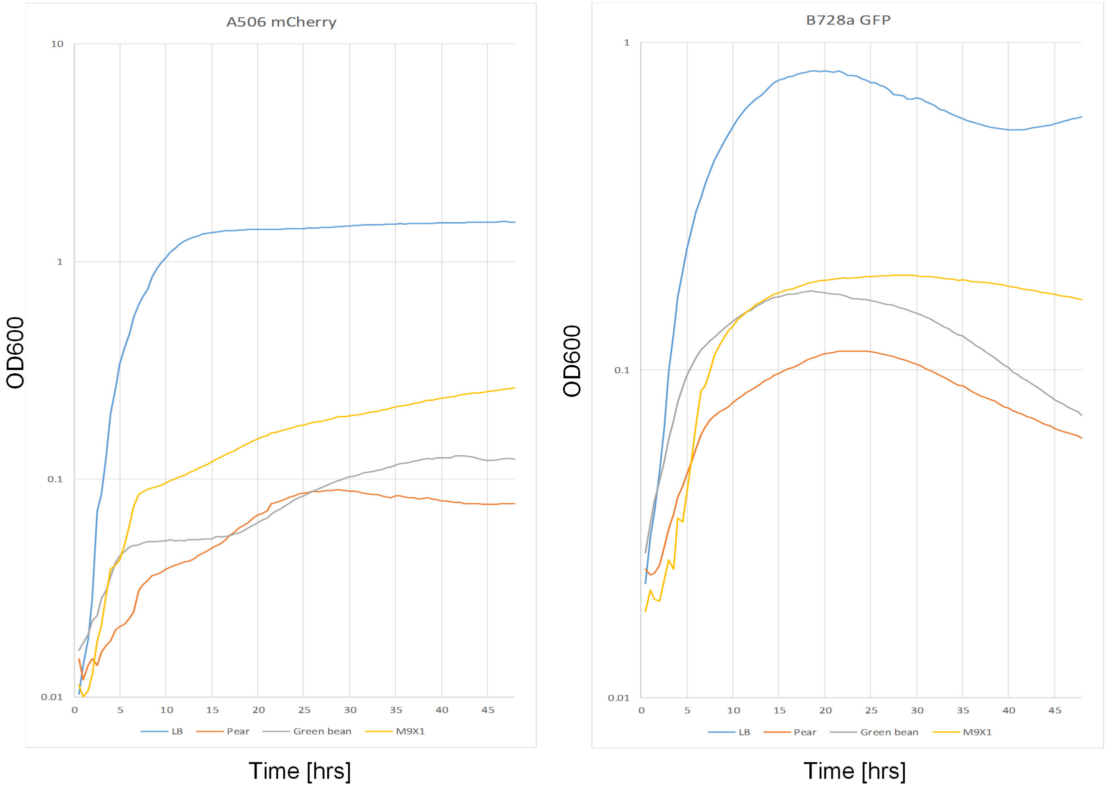


**Fig. S4 Growth curves of *P. fluorescens* A506 and *P. syringae* B728a grown in pear and green bean leaf solution.** *P. fluorescens* A506 mCherry and *P. syringae* B728a GFP were grown overnight in LB supplemented with gentamicin 30 µg/ml or kanamycin 50 µg/ml, respectively, at 28°C, 300 RPM. 3 ml LB with correlating antibiotics was inoculated with 100 µl of the overnight stock and grown until OD ~1.0 (in 1 cm cuvette). The bacteria were centrifuged, suspended in PBS 1X, diluted to OD 0.5 and diluted 1:20 once loaded into the 96-well plate. In order to mirror the growth conditions of the microscope experiment, both leaf wash solutions were diluted with PBS in the 96-well plate to reach the ratio of 1:5 PBS 1X and 4:5 LW. Triplicates for each condition and strain were used.

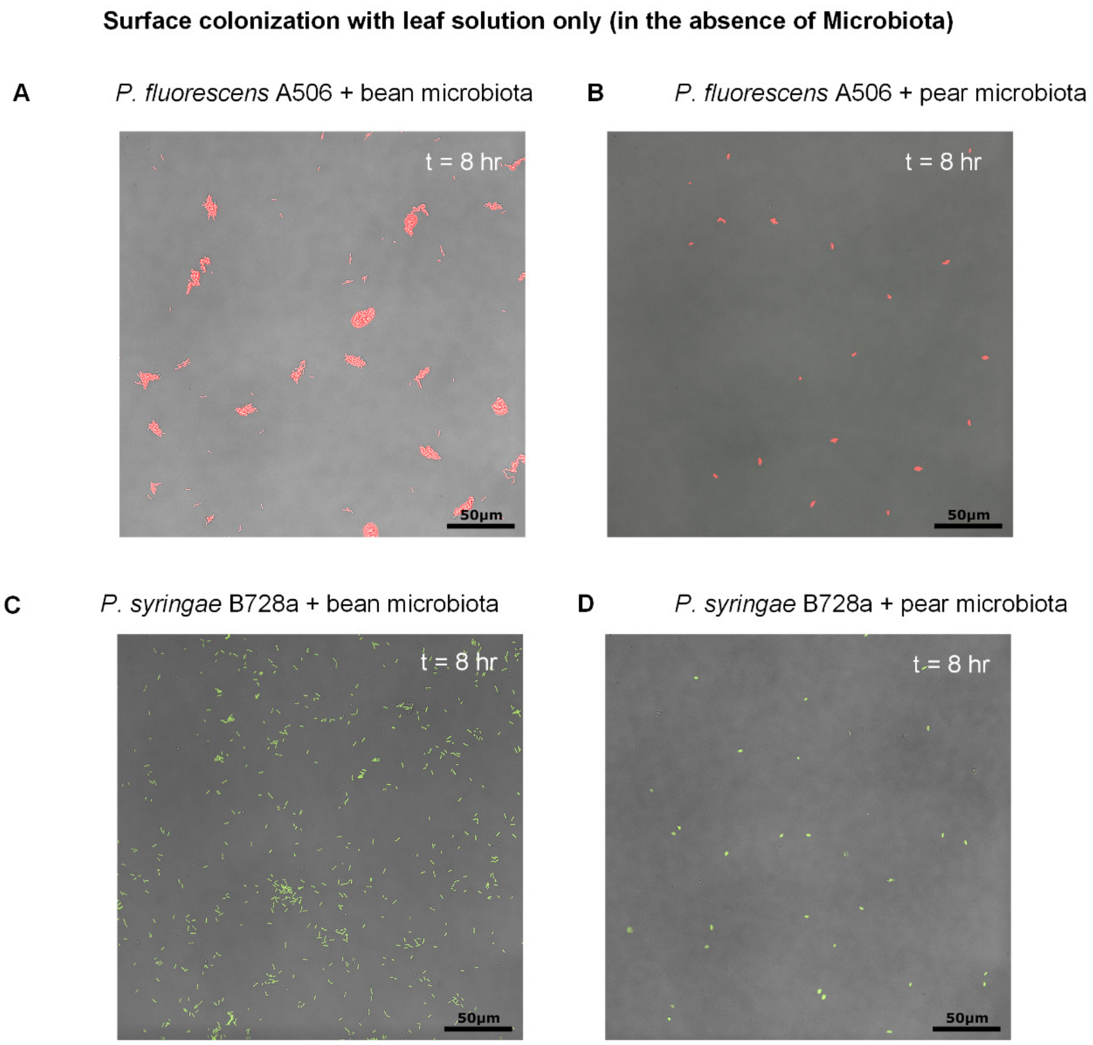


**Fig. S5. Surface colonization with leaf solution only (without microbiota).** Note the difference in colonization patterns: *Pf* A506 forms aggregates, while *Ps* B728a remains mostly as solitary cells that colonize the surface in a less ‘clustered’ manner.

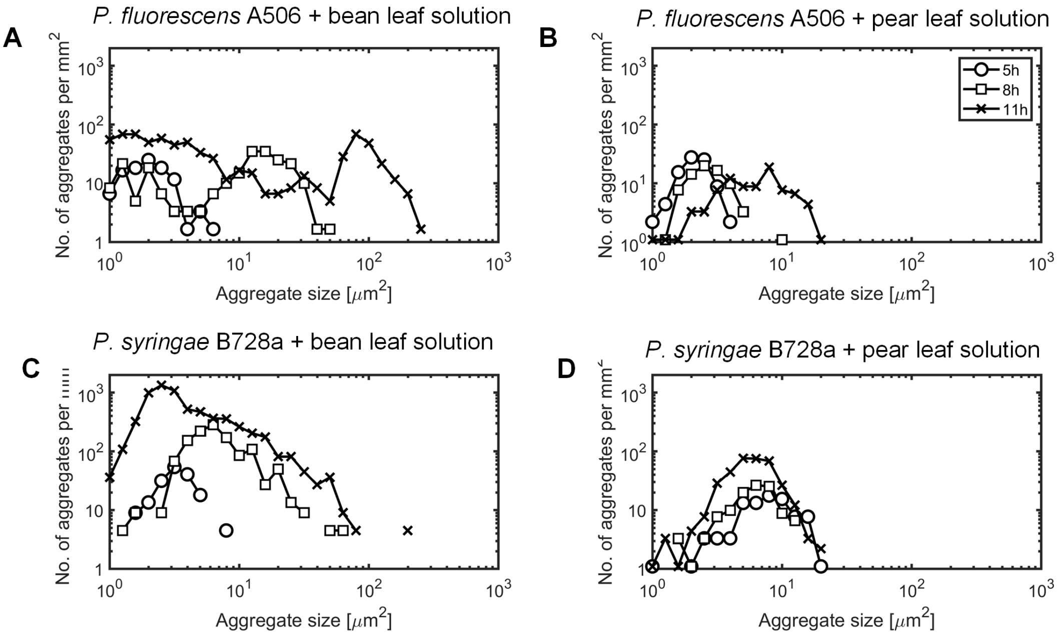


**Fig. S6 Aggregate-size distributions of immigrant bacteria in leaf solution (in the absence of microbiota).** Note marked growth either within fixed number of existing aggregates (*Pf* A506), or as larger number of dispersed individual cells (*Ps* B728a), **A-D.** Immigrant bacteria cluster-size distributions is given by area.

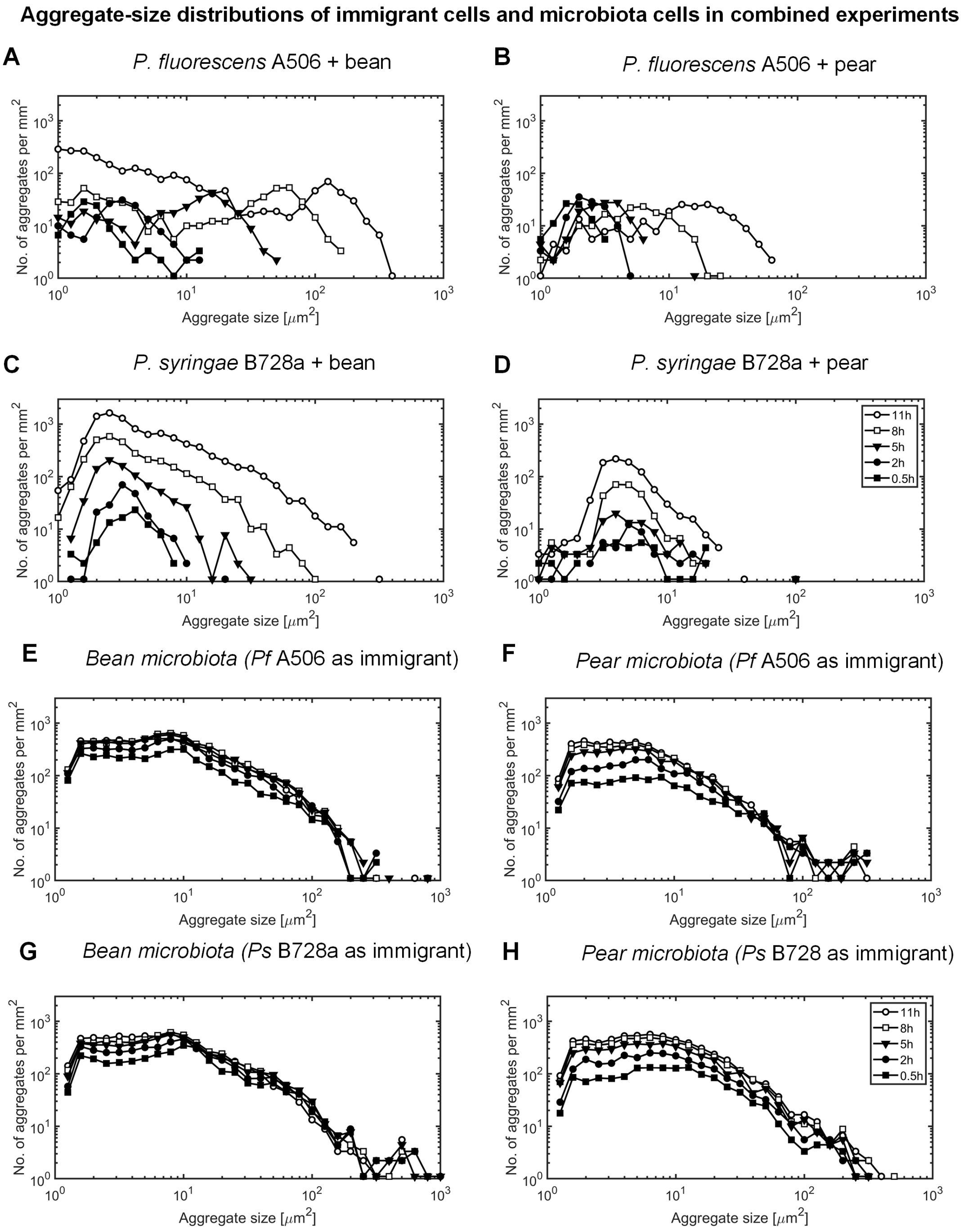


**Fig. S7 Aggregate-size distributions of immigrant bacteria and resident microbiota. A-D.** Immigrant bacteria aggregates (in presence of resident microbiota). **E-H.** Resident microbiota aggregates (in the presence of immigrant bacteria). The five lines represent different times: 0.5, 2, 5, 8, 11 hrs post inoculation.

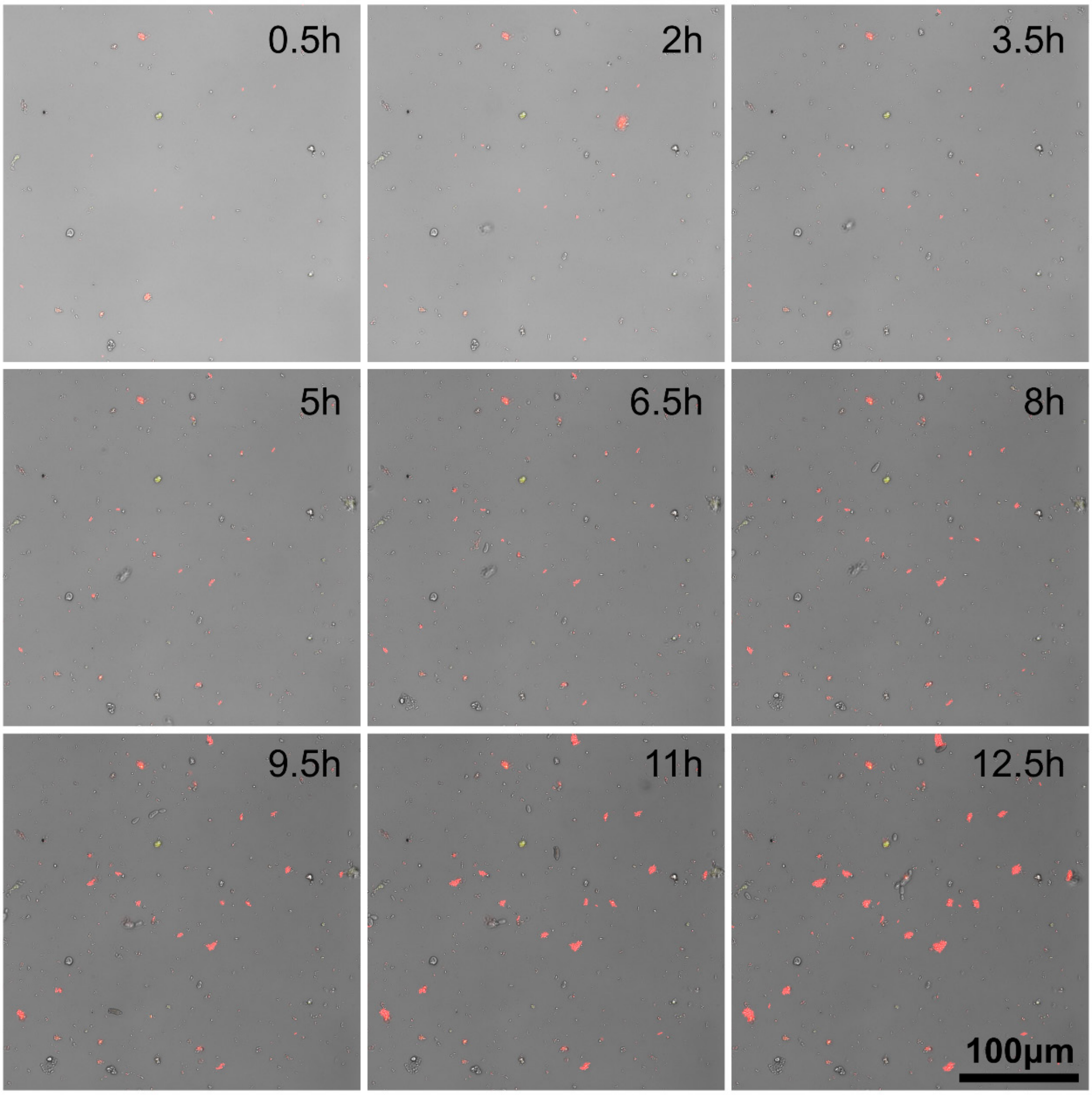


**Fig. S8. Time series of surface colonization of *Pf* A506 (red) with pear microbiota (Bright-field).** A single field of view is shown (out of the 9 field of views analyzed in the Results).

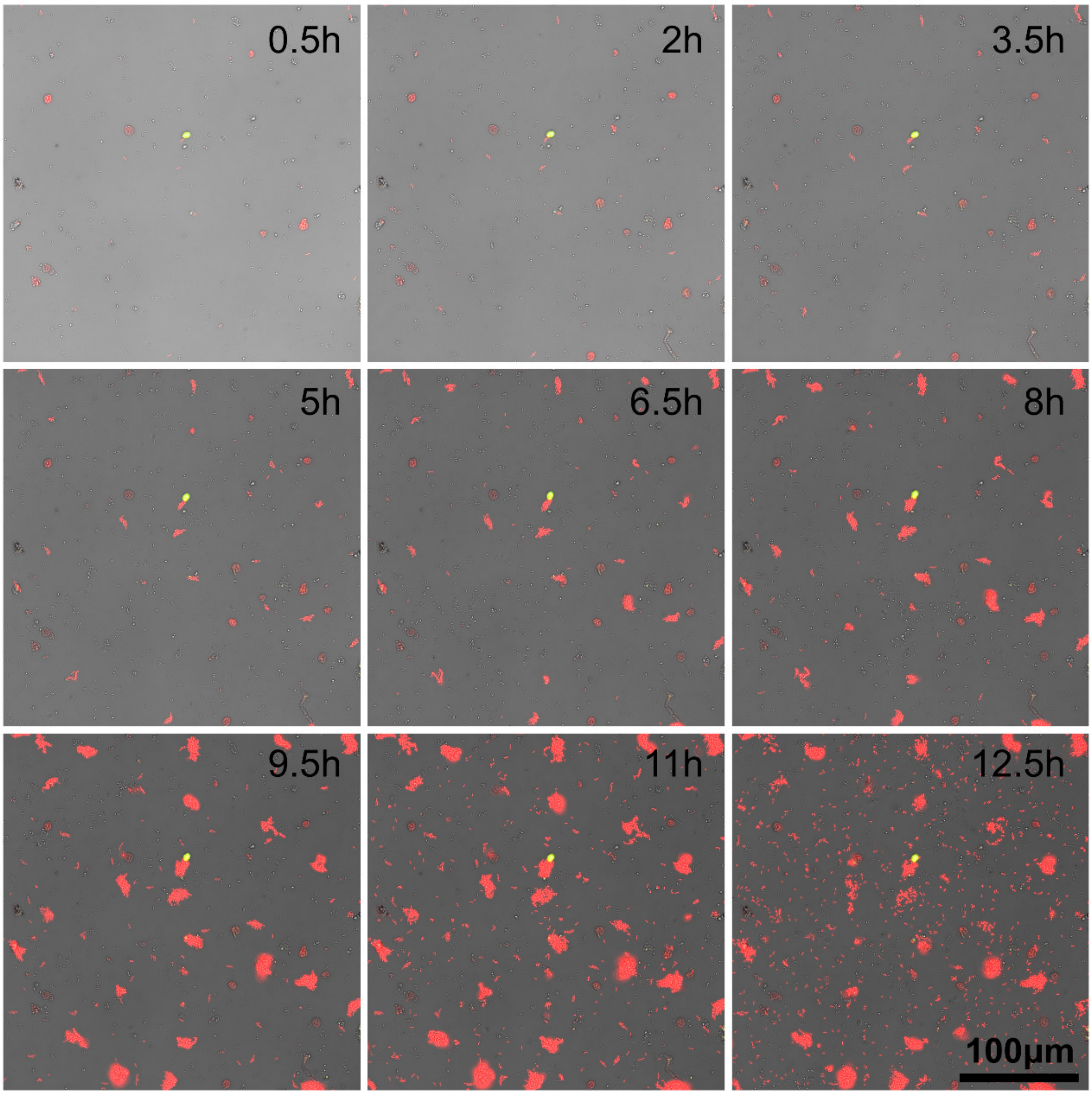


**Fig. S9. Time series of surface colonization of *Pf* A506 (red) with bean microbiota (Bright-field).** A single field of view is shown (out of the 9 field of views analyzed in the Results).

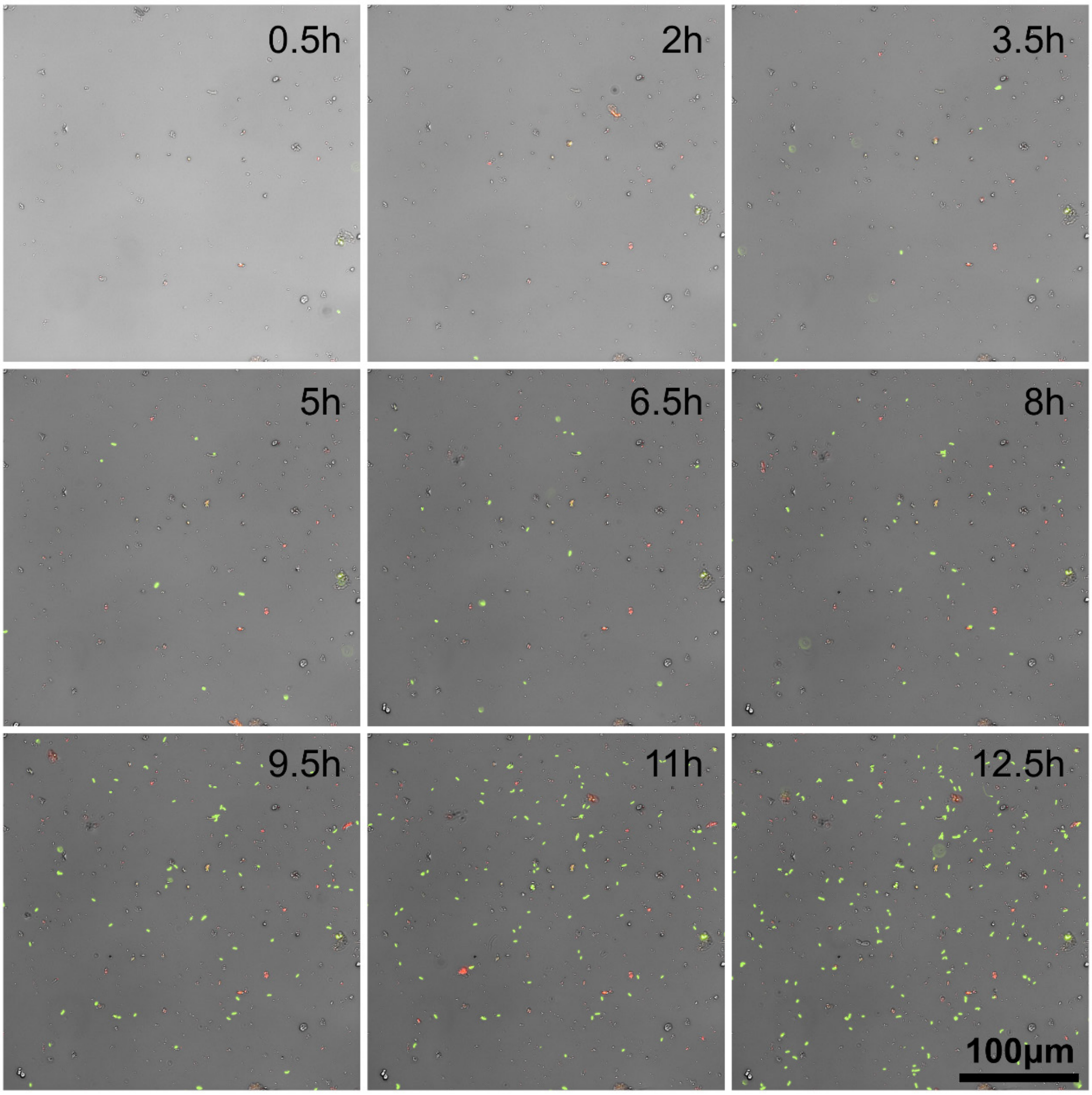


**Fig. S10. Time series of surface colonization of *Ps* B728a (green) with pear microbiota (Bright-field).** A single field of view is shown (out of the 9 field of views analyzed in the Results). Some plant derived particles such as chloroplasts can be observed as well (these were removed from the spatial analysis, see Methods and Fig. S2).

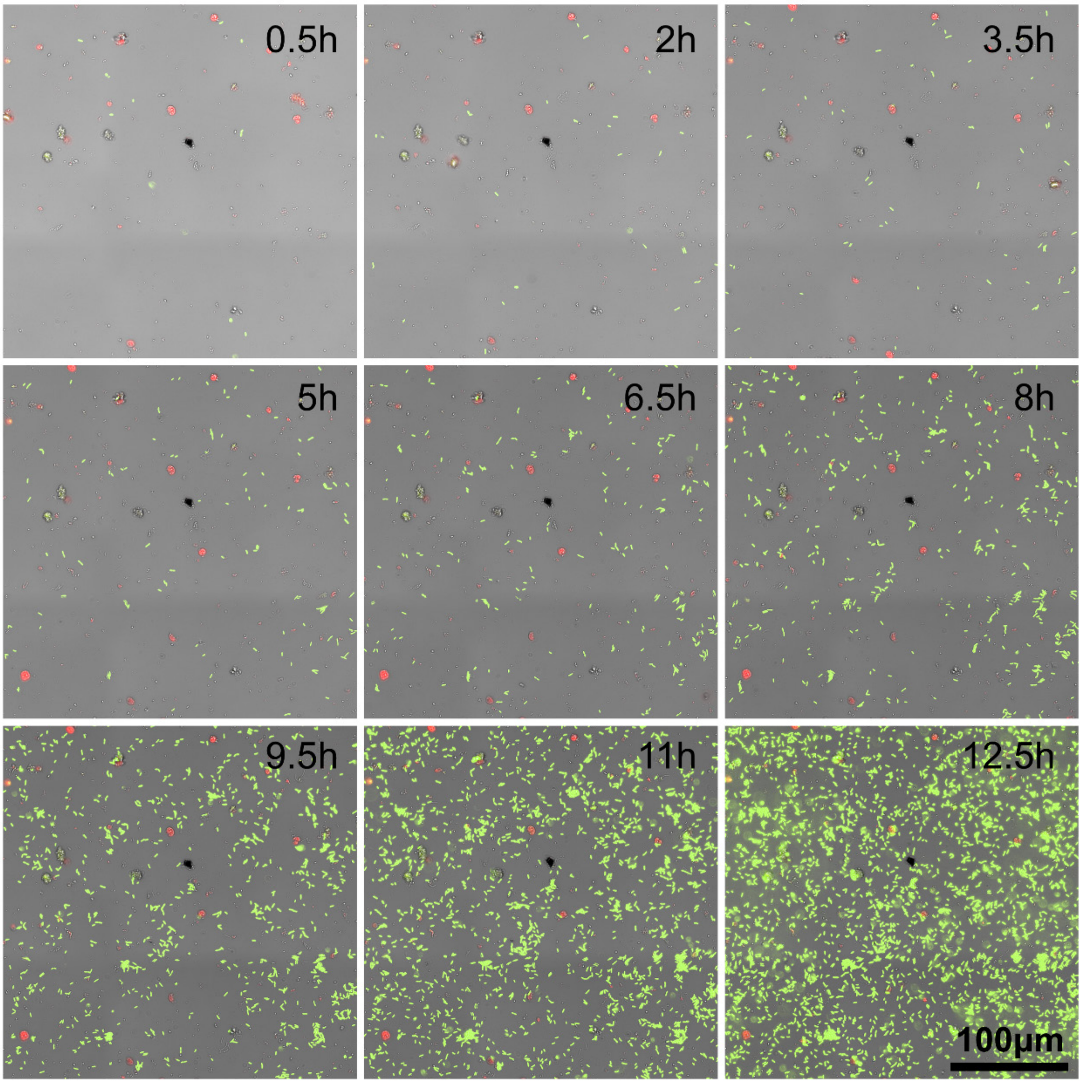


**Fig. S11. Time series of surface colonization of *Ps* B728a (green) with bean microbiota (Bright-field).** A single field of view is shown (out of the 9 field of views analyzed in the Results). Some plant particles and microbiota cells show weak autofluorescence observed in the green and red channels (these cells can be distinguished from immigrant cells as described in Methods and Fig. S2).

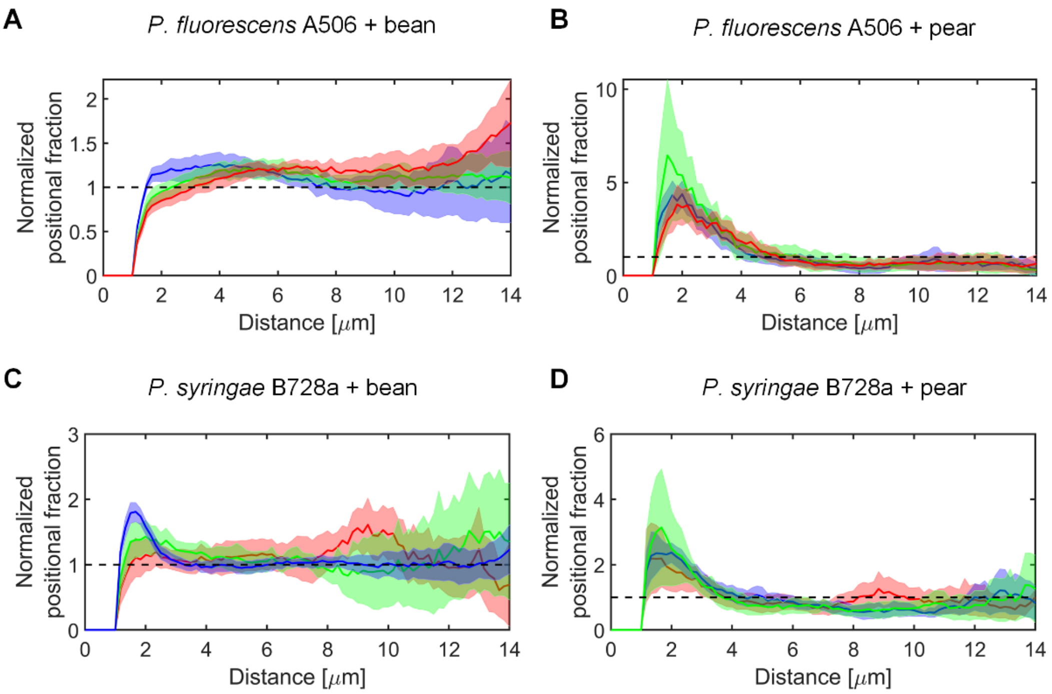


**Fig. S12 Nearest-neighbor analysis between immigrant cells and microbiota of three independent experiments.** Results are shown in blue, green, and red for three replicates (independent wells) of each of the four immigrant-microbiota pairs. The mean and 95% confidence intervals are based on pooling the results of 9 tiles per field of view per well (see Methods). The results shown in Fig. 5 in main text are shown here in blue. We note that for *Ps. syringae* B728a + bean (panel C), the area of the surface section analyzed in the two additional wells (red and green) was smaller than the area analyzed in the first well (blue), which may be related to the weaker deviation of the observed distribution from random.

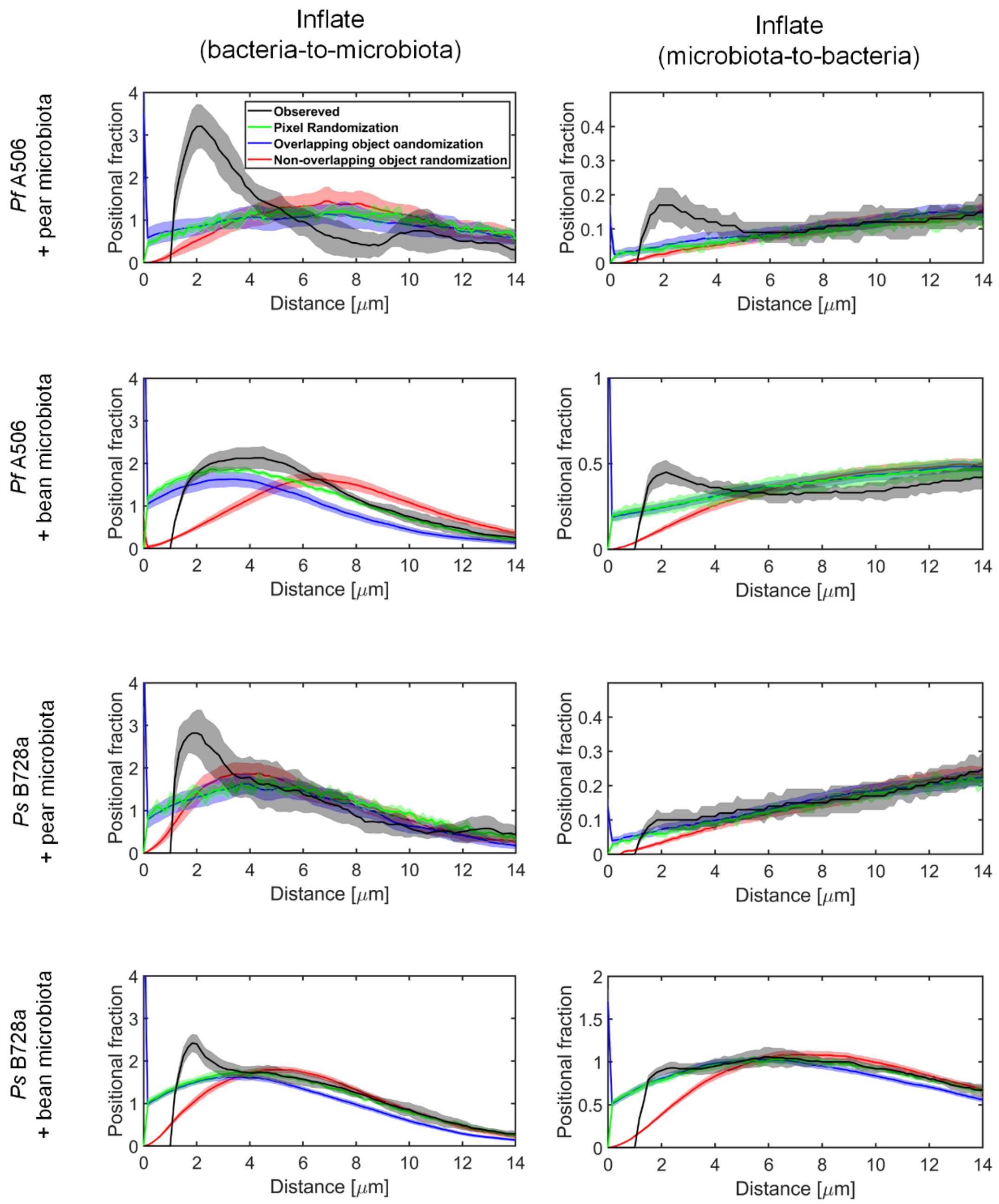


**Fig. S13 Comparing observed and various randomization schemes based on Monte-Carlo simulations.** The observed, non-normalized nearest-neighbor distribution (in black) is compared to per-pixel randomization (equivalent to CSR) (in green), and per-object Monte-Carlo randomization with and without allowing overlaps (in blue and red, respectively). Observed and per-pixel randomization mean and 95% CI based on pooling 9 tiles per well, object based randomizations mean and 95% CI based on 5 randomizations for each of the 9 tiles (45 randomizations in total per well). Generally, the deviation from random of the observed data is larger if normalized by the non-overlapping-objects model, indicating that co-localization of immigrant cells and native microbiota might be under-stated under the assumption of CSR (as shown in Figure 5).

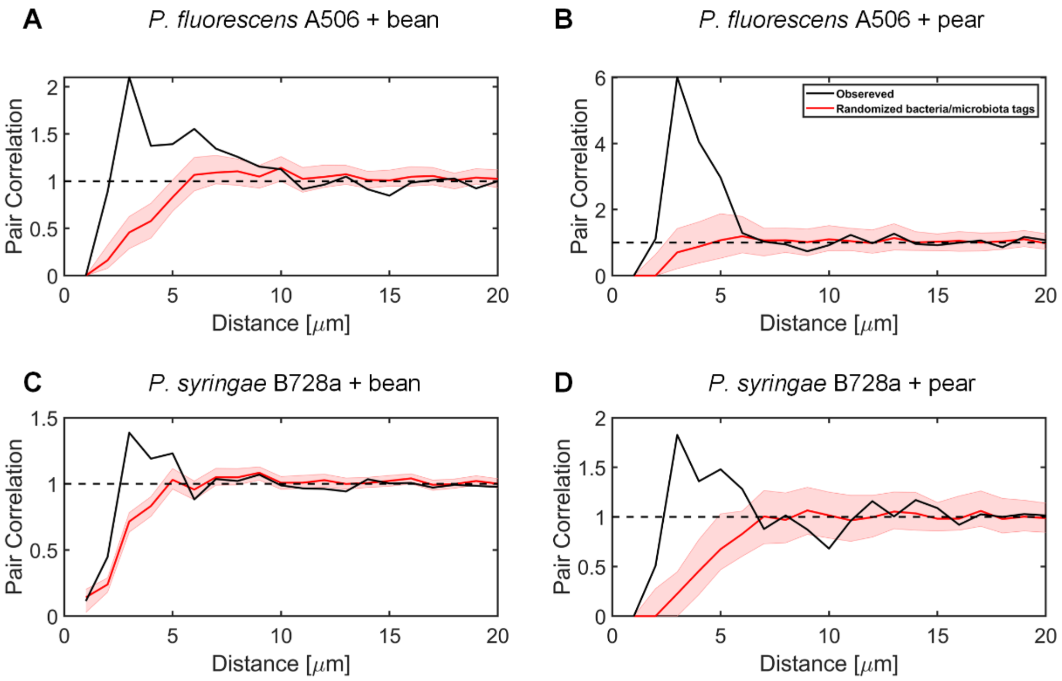


**Fig. S14 Randomization by permutations of immigrant/microbiota tags.**

Results are by a center-to-center PCC method. The observed PCC analysis (in black) is based on the entire field of view. The expected mean and 95% CI (in red) are based on PCC analysis of 100 random permutations of the class tags of immigrant and microbiota objects.

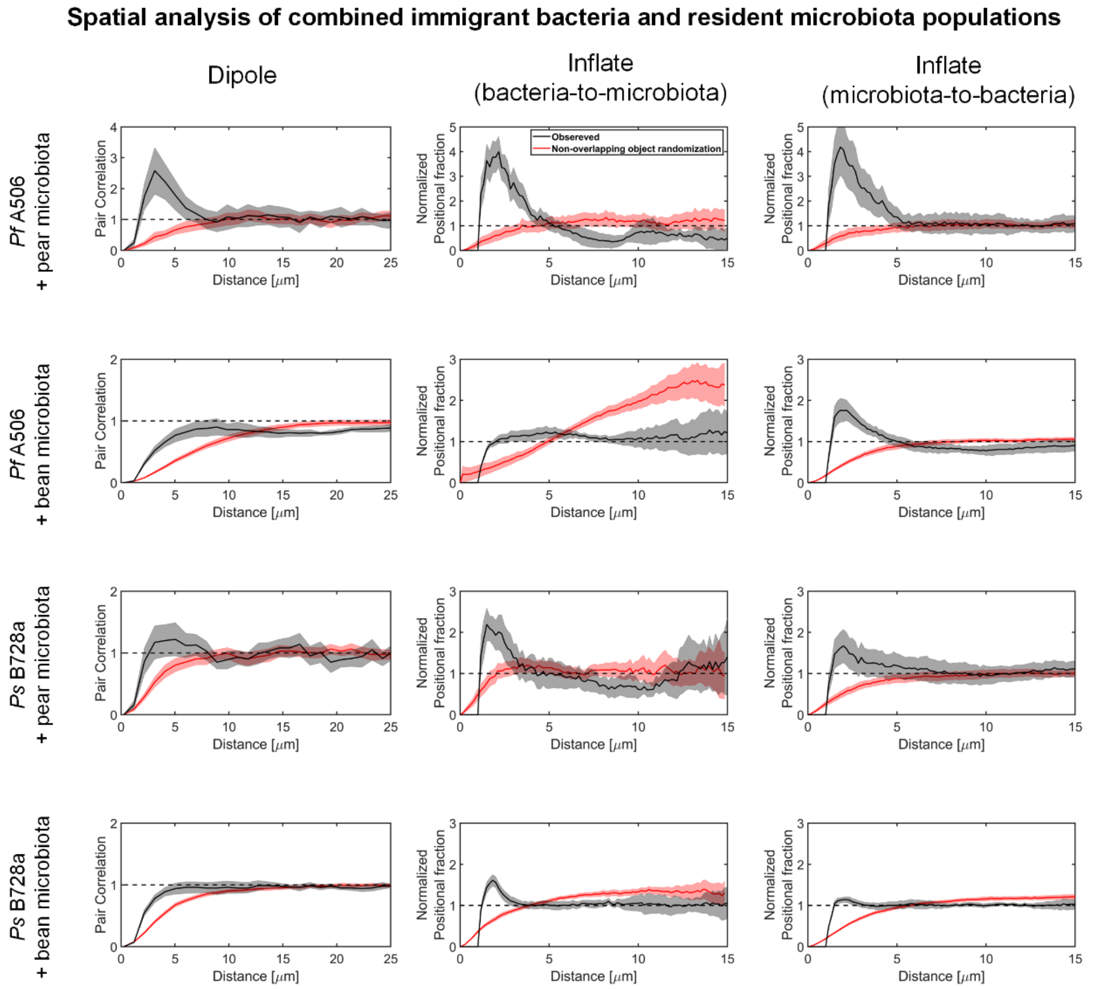


**Fig. S15 Comparing observed and Monte-Carlo simulations of immigrant bacteria vs. native microbiota.** Data is as in Fig. 5 (Main text), with the addition of the per-object Monte-Carlo randomization without allowing overlaps (in red). Observed data envelopes (gray) represent 95% CI based on pooling 9 tiles per well, object based randomizations mean and 95% CI based on 5 randomizations for each of the 9 tiles (45 randomizations in total per well). Generally, the deviation from random of the observed data is larger if compared to the non-overlapping-object randomization model, indicating that co-localization of immigrant cells and native microbiota might be under-stated under the assumption of CSR (as shown in Figure 5).

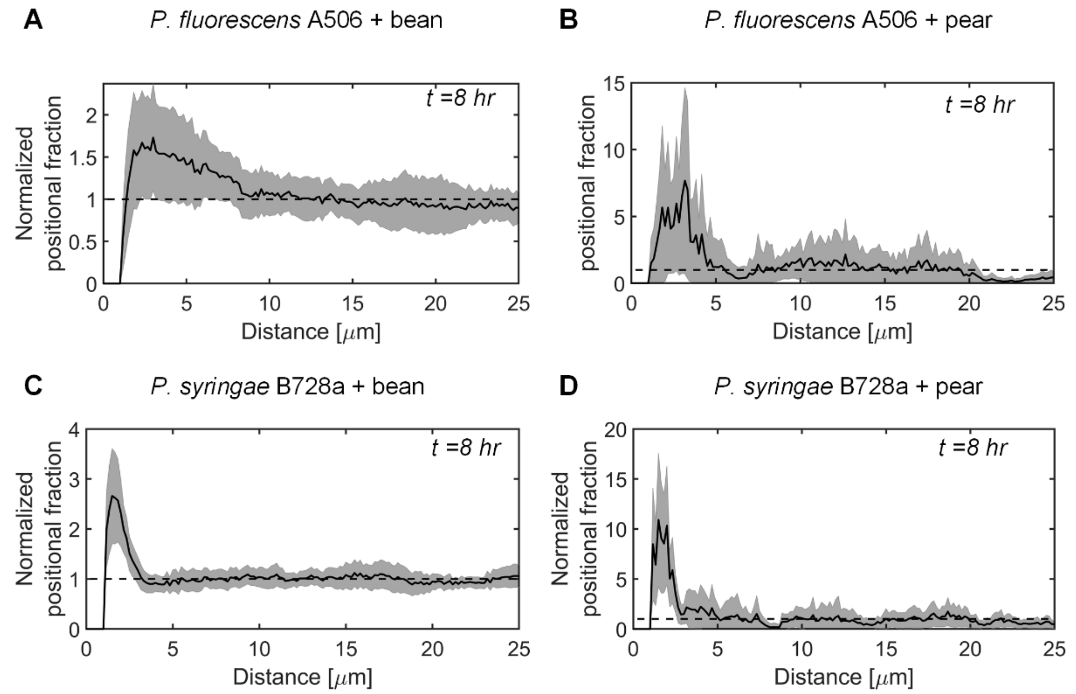


**Fig. S16 Spatial analysis of bacteria vs. plant-derived-particles.** Nearest Neighbor analysis carried out by 2D inflate analysis, using DAIME. Mean+95% CI of 9 tiles per well. Distances are measured between bacterial structures and plant parts (see classifications at Supp. Fig. S2). Though limited by a small dataset (plant parts were between 0.4% to 3.9% of total number of objects, see Supp. Table S1), the analysis indicates that *Pf* B728a shows some attraction to plant parts, roughly equivalent to its attraction to native microbiota. *Ps* A506 show less significant affinity to plant parts.

**Supp. Table. S1** Statistics of particle classification (classification was done manually by two of the co-authors).

| | | By area | | By objects | |
|---|---|---|---|---|---|
| | | Microbiota | Plant | Microbiota | Plant |
| Classification I: | a506+pear | 95.9% | 4.1% | 99.3% | 0.7% |
| | b728+pear | 97.0% | 3.0% | 99.6% | 0.4% |
| | a506+bean | 91.7% | 8.3% | 98.7% | 1.3% |
| | b728+bean | 90.0% | 10.0% | 98.0% | 2.0% |
| Classification II: | a506+pear | 92.0% | 8.0% | 97.9% | 2.1% |
| | b728+pear | 91.6% | 8.4% | 98.6% | 1.4% |
| | a506+bean | 84.8% | 15.2% | 97.0% | 3.0% |
| | b728+bean | 90.6% | 9.4% | 96.1% | 3.9% |

**Additional Supplementary files**

**File S1 Two-way microscale interaction dynamics: *Pf* A506 + pear microbiota.** Example time-series images in the microscopic neighborhood of microbiota and immigrant cell clusters.

**Fig. S2 Two-way microscale interaction dynamics. *Pf* A506 + bean microbiota.** Example time-series images in the microscopic neighborhood of microbiota and immigrant cell clusters.

**Fig. S3 Two-way microscale interaction dynamics. *Ps* B728a + pear microbiota** Example time-series images in the microscopic neighborhood of microbiota and immigrant cell clusters.

**Fig. S4 Two-way microscale interaction dynamics. *Ps* B728a + bean microbiota** Example time-series images in the microscopic neighborhood of microbiota and immigrant cell clusters.

**Fig. S5**. **Raw data for spatial analysis** (zip file). Segmentation masks of immigrant bacteria and native microbiota at t=8hr. 5960x5960 pixels at 0.16 um/pixel. CCITT RLE compression, two color .tiff format.
